# ImuA participates in SOS mutagenesis by interacting with RecA1 and ImuB in *Myxococcus xanthus*

**DOI:** 10.1101/2020.12.14.422803

**Authors:** Duohong Sheng, Ye Wang, Zhiwei Jiang, Dongkai Liu, Yuezhong Li

## Abstract

Bacteria have two pathways to restart stalled replication forks caused by environmental stresses, error-prone translesion DNA synthesis (TLS) catalyzed by TLS polymerase and error-free template switching catalyzed by RecA, and their competition on the arrested fork affects bacterial SOS mutagenesis. DnaE2 is an error-prone TLS polymerase, and its functions require ImuA and ImuB. Here we investigated the function of *imuA*, *imuB* and *dnaE2* in *Myxococcus xanthus* and found that *imuA* showed differences from *imuB* and *dnaE2* in bacterial growth, resistance and mutation frequency. Transcriptomics analysis found that ImuA were associated with bacterial SOS response. Yeast-two-hybrid scanning revealed that ImuA interacted with RecA1 besides ImuB. Protein activity analysis proved that ImuA had no DNA binding activity, but inhibited the DNA binding and recombinase activity of RecA1. These findings highlight that ImuA not only participates in TLS by binding ImuB, but also inhibits the recombinase activity of RecA1 in *M. xanthus*, suggesting a role of ImuA in the two replication restart pathways.

**Importance:** DnaE2 is responsible for bacterial SOS mutagenesis in nearly one third of sequenced bacterial strains. However, its mechanism, especially the function of its accessory protein ImuA, is still unclear. Here we reported that *M. xanthus* ImuA might facilitate DnaE2 TLS by inhibiting the recombinase activity of RecA1, which helps to explain the mechanism of DnaE2-dependent TLS and the scientific problem of choosing one of the two restart pathways to repair the stalled replication fork.

## Introduction

DNA damage caused by environmental stress might stall DNA replication, resulting in collapses of the replication fork and breaks in DNA strands, even cell death. Bacteria possess two pathways (also named postreplication repair pathways in some publications) to reinitiate the stalled replication fork: translesion DNA synthesis (TLS) and template switching (**Fig. S1**). Template switching is mediated by RecA-based homologous recombination and is error-free, while TLS is inherently error-prone, which bypasses the DNA lesion by inserting a nucleotide opposite of the lesion. TLS is the main source of SOS mutagenesis, which opportunistically leads to adaptive evolution (Agashe, 2017; Hershberg, 2015; Lovett, 2017; Shee et al., 2011).

*E. coli* contains three different TLS polymerases, Pol II (*polB*), Pol IV (*dinB*) and Pol V (*umuCD*), which are induced by the SOS response (Agashe, 2017; Goodman & Woodgate, 2013; Schlacher & Goodman, 2007). Compared with Pol II and Pol IV, Pol V catalyzes DNA synthesis with the lowest fidelity and is the main enzyme for bacterial SOS-induced mutations (Fuchs et al., 2004; Goodman & Woodgate, 2013; Jaszczur et al., 2016). However, only approximately one third (1,707 strains) of the sequenced bacteria (5,360 strains, KEGG data in Jan 2020) encodes Pol V. In those strains without Pol V, nearly a half (1,699 strains) contains DnaE2 (ImuC in some publications) protein that has been shown to play a key role in SOS mutagenesis (Abella et al., 2004; Alves et al., 2017; Baker & Kornberg, 1998; Boshoff et al., 2003; Erill et al., 2006; Galhardo et al., 2005; Tsai et al., 2012; Timinskas et al., 2014; Warner et al., 2010). DnaE2 is a structural homolog of DnaE1 subunit of replicative DNA polymerase III. Unlike DnaE1, DnaE2 does not bind to the *ε* subunit (3’ to 5’ exonuclease activity) of DNA polymerase III, and the DNA synthesis catalyzed by DnaE2 has no proofreading function, resulting in a high mutation frequency. Additionally, DnaE2 lacks the C-terminal domain (CTD) to bind to the τ subunit which acts as a loading subunit to assemble DnaE onto the replication fork (Boshoff et al., 2003; Timinskas et al., 2014). Thus, without aids from other cofactors DnaE2 does not function in the replication of chromosomal DNA.

The *imuA* and *imuB* genes, usually located in the SOS-regulated operon of *imuA-imuB-dnaE2*, are required for DnaE2 mutagenesis (Abella et al., 2004; Alves et al., 2017; Erill et al., 2006; Galhardo et al., 2005). The *imuA* or *imuB* mutants of *Caulobacter crescentus* and *Mycobacterium tuberculosis* showed a significant reduction in SOS-induced mutation frequency, similar to that of the *dnaE2* mutants (Galhardo et al., 2005; Warner et al., 2010). ImuB is the member of UmuC subfamily of the Y-family DNA polymerases, lacks three amino acid residues in the active center and probably has no DNA polymerase activity. ImuB retains a beta-clamp-binding motif (^354^QLPLWG^359^), and thus is able to interact with β-clamp. Warner et al. speculated that the interaction between ImuB and β-clamp mediates the access of DnaE2 to the replication fork, leading to DnaE2-induced mutation (Warner et al., 2010).

ImuA is a RecA-like protein that contains the conserved ATPase core domain but lacks the typical N- and C-termini of RecA. RecA was reported to be involved in the error-prone TLS pathway catalyzed by Pol V via active mutasome formation (Pol V-RecA-ATP) (Jiang et al., 2009). However, RecA was not involved in DnaE2 mediated error-prone TLS pathway (Alves et al., 2017). Similarly, ImuA did not interact directly with DnaE2 (Warner et al., 2010). Thus, the role of ImuA in DnaE2 mediated TLS does not imitate the PolV-RecA’s action and the ImuA’s function is still unclear.

*Myxococcus xanthus* DK1622 is the model strain of myxobacteria, and its genome size is greater than 9 Mbp with a large number of DNA repeats, such as duplicate *recA* genes (Chen et al., 2016; Goldman et al., 2006). Recently, we showed that the expression of RecA1 (MXAN_1441) with recombinase activity is very low in *M. xanthus* DK1622 cells (Sheng et al., 2020), which might limit the template switching events at stalled forks. Our previous studies have showed that *M. xanthus dnaE2* gene (*MXAN_3982*) encoded an error-prone DNA polymerase for DNA replication. The absence of *dnaE2* showed a significant decrease in the mutation frequency (Peng et al., 2017). These above results suggest that *M. xanthus* is an ideal strain for studying DnaE2 mutagenesis, because the recombination-mediated template switching is weak, while the DnaE2-catalysed TLS pathway plays a more important role in restarting replication forks in *M. xanthus* DK1622. Herein, we further showed that ImuA probably functions as a mediator to balance the template switching and TLS pathways via its competitive binding capacity to RecA1 and ImuB.

## Results

### 1. Induction of *imuA* by UV irradiation occurs earlier than that of *imuB* and *dnaE2*

The *imuA*, *imuB,* and *dnaE2* genes, adjacently located either in an operon or not, are SOS-regulated genes and can be induced by DNA-damaging agents, such as UV light. (Abella et al., 2004; Erill et al., 2006; Galhardo et al., 2005; Warner et al., 2010). In *M. xanthus* DK1622, these three genes are arranged in an A-B-E2 order; *imuA* and *imuB* are adjacent, and *dnaE2* is located downstream, separated from the two genes by a 6.57-kb fragment containing seven cistrons (**Fig. 1A**). The three genes each possess independent promoter region, and thus transcribe independently (**Fig. 1B**). There is a delta-proteobacterial SOS box sequence (CTRHAMRYBYGTTCAGS) in the promoter region of *dnaE2* or *imuB*, but not *imuA*.

**Figure 1.**
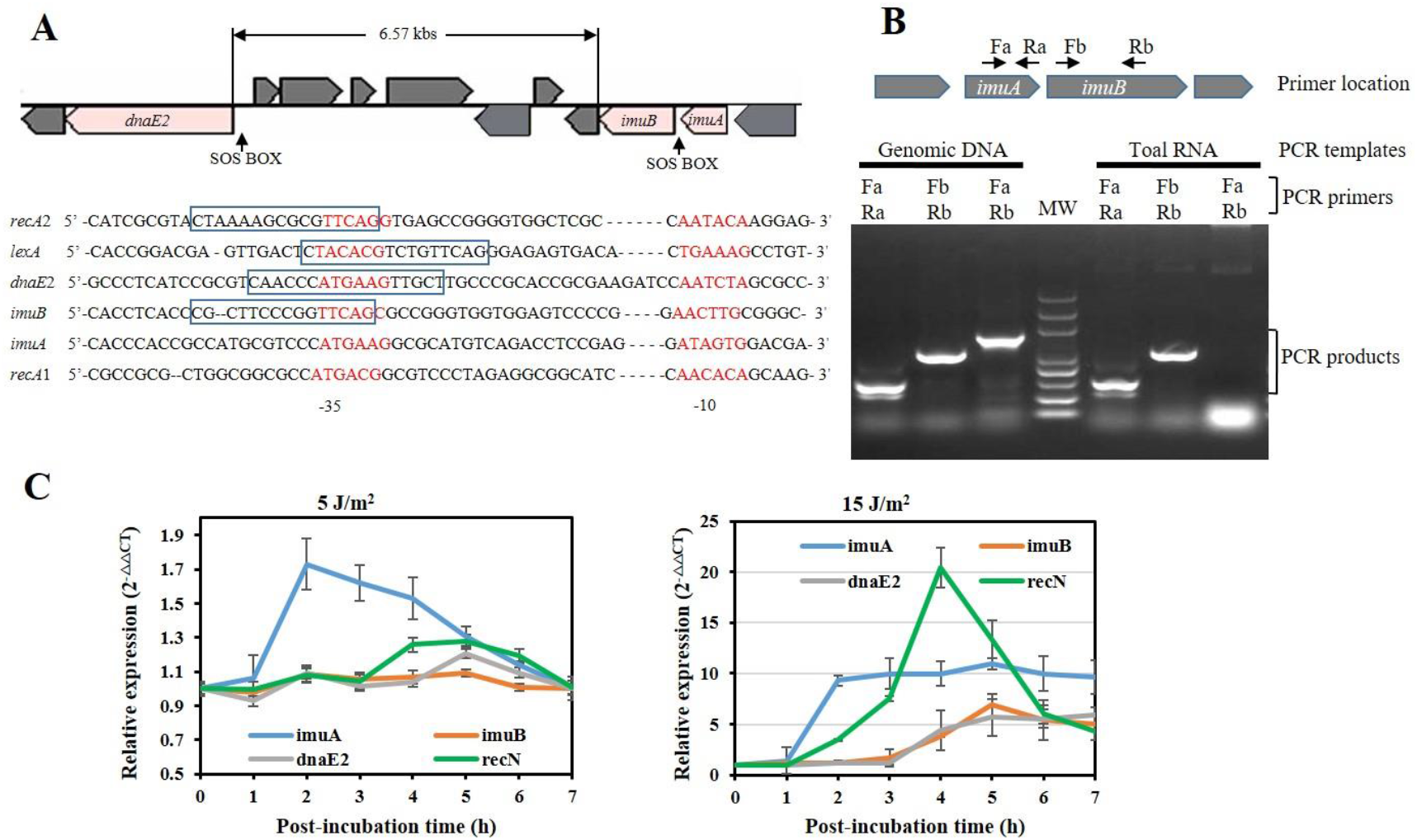
Gene location and transcription analysis of *imuA, imuB* and *dnaE2* in *M. xanthus* DK1622. (A) Schematic gene location and the promoter alignment. The RNA polymerase binding regions (−10 and −35) are marked in red and the SOS box regions are in blue frame. The promoters of *recA1, recA2* and *lexA* are also provided for comparison. (B) PCR amplification of *imuA-imuB* intergenic region. Primers used for the PCR amplification are indicated above the gel map. Total RNA was extracted from *M. xanthus* and reverse transcribed into cDNA for PCR. Genomic DNA was used for control. (C) Expression induction analysis of *imuA, imuB* and *dnaE2* with UV irradiation. The DK1622 cells were exposed to UV irradiation at the dose of 5 or 15 J/m^2^ using a UV crosslinking machine. After irradiation, the samples were post-incubated at 30 °C for different h (from 0 to 7 h) and then the total RNA were extracted from cell for RT-PCR. The relative gene expression quantified by the comparative CT (2^−ΔΔCT^) method. Error bars represent means ± SEM (n=3, p<0.05).

In UV-irradiated *M. xanthus* cells, the expression levels of *imuA*, *imuB,* and *dnaE2* genes were induced with significantly different induction times and strengths (**Fig. 1C**). Under the low-dose UV radiation treatment (5 J/m^2^), the *imuA* transcription was upregulated 1 h after irradiation, *dnaE2* responded to irradiation 2 h later, and *imuB* was not induced. When treated with a high-dose radiation, the transcriptions of *imuA*, *imuB* and *dnaE2* were all significantly upregulated: *imuA* was upregulated 1 h after irradiation, while *imuB* and *dnaE2* were 2 h later. Notably, *imuA* showed a higher transcription rate than that of *imuB* and *dnaE2* in response to a high dose of UV irradiation.

### 2. Compared with *imuB* or *dnaE2* mutant, *imuA* mutant reduced the UV sensitivity on the growth, mutation and survival of *M. xanthus*

To investigate the functions of the three genes, we constructed deletion mutants of *imuA* and *imuB* (named IA and IB) (**Fig. S2**). The deletion of *dnaE2* (YL1601) (Peng et al., 2017), *imuB* or *recA1* (Sheng et al., 2020) did not significantly affect cellular growth, but the *imuA* mutant showed approximate 12-h delay before the exponential growth stage though its maximum growth rate (μ_max_) did not change significantly. UV irradiation inhibited the growth of DK1622, YL1601 (Δ*dnaE2*), and IB (Δ*imuB*), whereas the growth curve of IA (Δ*imuA*) remained unaltered before and after irradiation (**Fig. 2**). It suggested that *imuA* might impede radiation-induced growth inhibition as the irradiated cells need more time to repair DNA damage (Aksenov, 1999; Hamkalo & Swenson, 1969).

**Figure 2.**
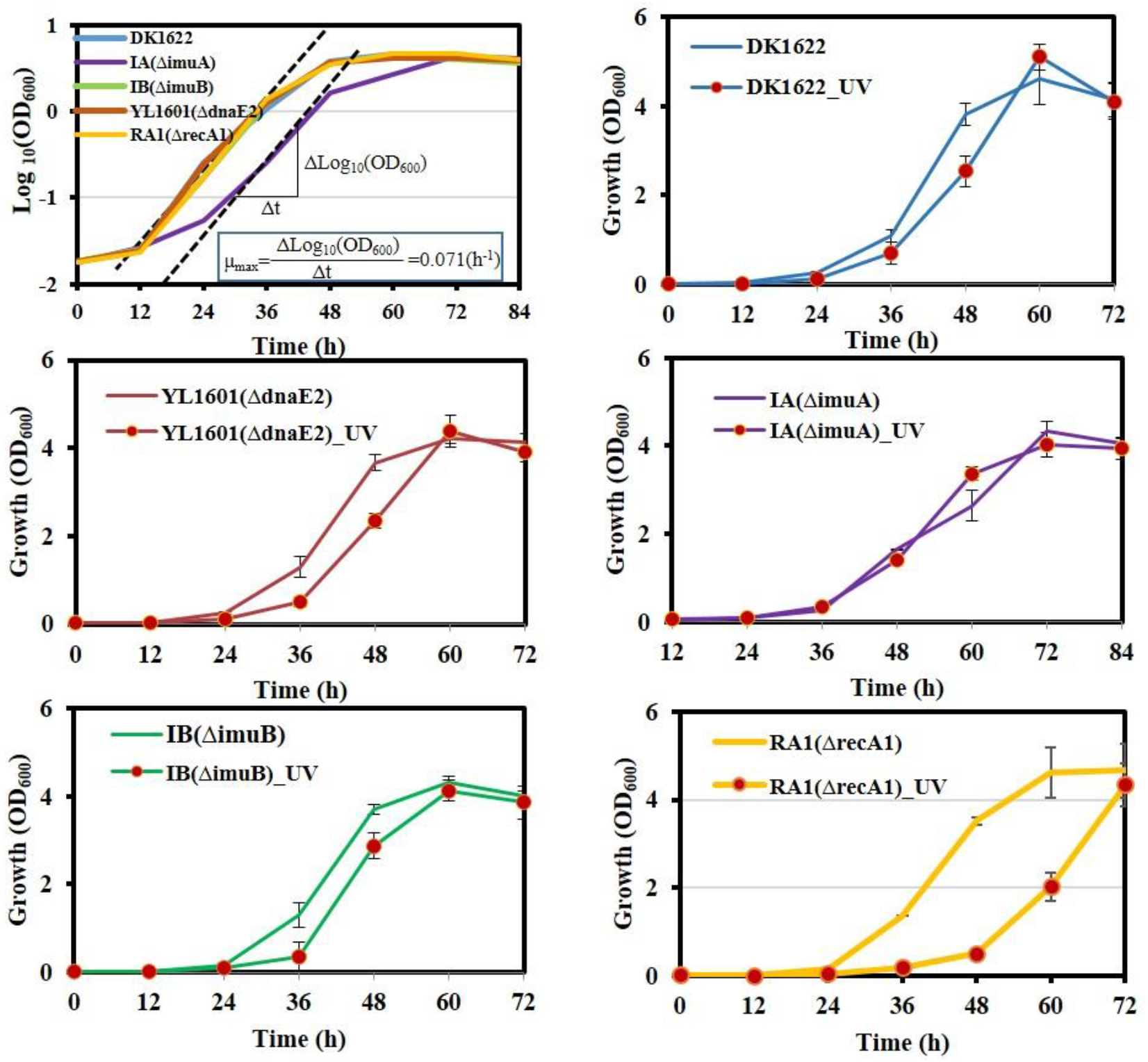
Growth analysis of the *imuA, imuB* and *dnaE2* mutants with and without UV irradiation, wild-type *M. xanthus* DK1622 and the mutant of SOS gene *recA1* as controls. Strains were treated with 15 J/m^2^ UV and then shaken at 30 °C for different time to measure the OD_600_ values of culture. DK, wild-type *M. xanthus* DK1622; IA, the *imuA* mutant; IB, the *imuB* mutant; YL1601, the *dnaE2* mutant. Error bar indicates the standard error of the mean of six independent experiments.

SOS mutagenesis was analyzed by calculating the target-site mutation frequency of nalidixic acid of *M. xanthus* in response to UV irradiation (Peng et al., 2017). In the presence of nalidixic acid and without UV irradiation, the mutation frequency of wild-type *M. xanthus* cells was approximately 3 × 10^−8^, in line with the previous report (Peng et al., 2017). Further UV irradiation doubled the mutation frequency of *M. xanthus* (**Fig. 3A**). Deletion of *dnaE2* or *imuB* significantly reduced the mutation frequency of this bacterium, whether it was radiation or not, which suggested that ImuB and DnaE2 were responsible for bacterial mutation frequency. Deletion of *imuA* also significantly decreased the mutation frequency, but it was not as obvious as that of *imuB* or *dnaE2*. In particular, the mutation frequency of the *imuA* mutant still retained UV inducibility (**Fig. 3A**), which suggested that ImuA also involved in the mutation but probably in a different manner from that of ImuB or DnaE2. The survival analysis against UV radiation showed that the absence of *imuA* slightly decreased the survival rate of *M. xanthus* cells with the increase of radiation dose, and the absence of *imuB* or *dnaE2* had a more significant effect (**Fig. 3B**). The results also proved that the functions of ImuA and ImuB (or DnaE2) were not synchronized.

**Figure 3.**
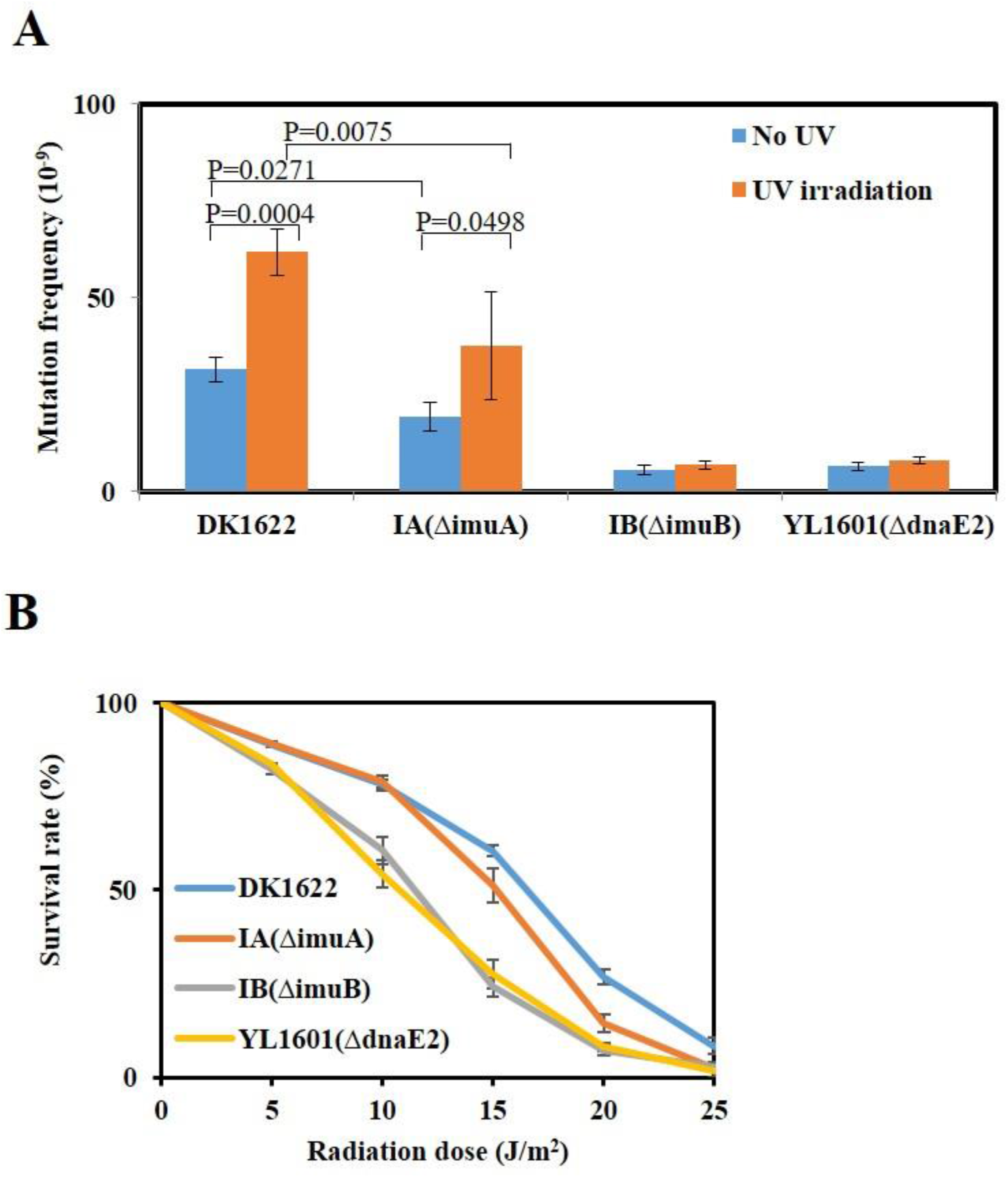
Mutability and survival analysis of the *imuA, imuB* and *dnaE2* mutants, compared with *M. xanthus* DK1622. (A) Mutability analysis. 5 × 10^9^ cells (irradiated with UV) were spread on CYE agar containing 40 μg/ml nalidixic acid and the resistant clones were counted to represent the mutation ability. Error bar indicates the standard error of the mean of four independent experiments. (B) Survival analysis with different doses of UV irradiation. DK, wild-type *M. xanthus* DK1622; IA, *imuA* mutant; IB, *imuB* mutant; YL1601, *dnaE2* mutant. Error bar indicates the standard error of six independent experiments.

### 3. ImuA affects the expression of genes related to the SOS response

To understand potential roles of *imuA*, transcriptomes of the *imuA* deleted and overexpressing strains were compared with that of the wild type *M. xanthus* DK1622 strain. We identified a total of 260 differentially expressed genes (DEGs), out of which 81 DEGs were inhibited by ImuA (upregulated in the *imuA* deletion mutant and downregulated in the *imuA* overexpression strain) and 179 were increased by ImuA (downregulated in the *imuA* deletion mutant and upregulated in the *imuA* overexpression strain) (**Tables S1** and **S2**). 135 of the 260 DEGs were known and functionally classified into genetic information processing (63 genes), metabolism (47 genes), environmental information processing (23 genes) and cellular processes (2 genes) (**Fig. 4A**). Notably, the majority of the genes belonging to genetic information processing were inhibited by ImuA, while most genes of metabolism and environment information processing were promoted by ImuA.

**Figure 4.**
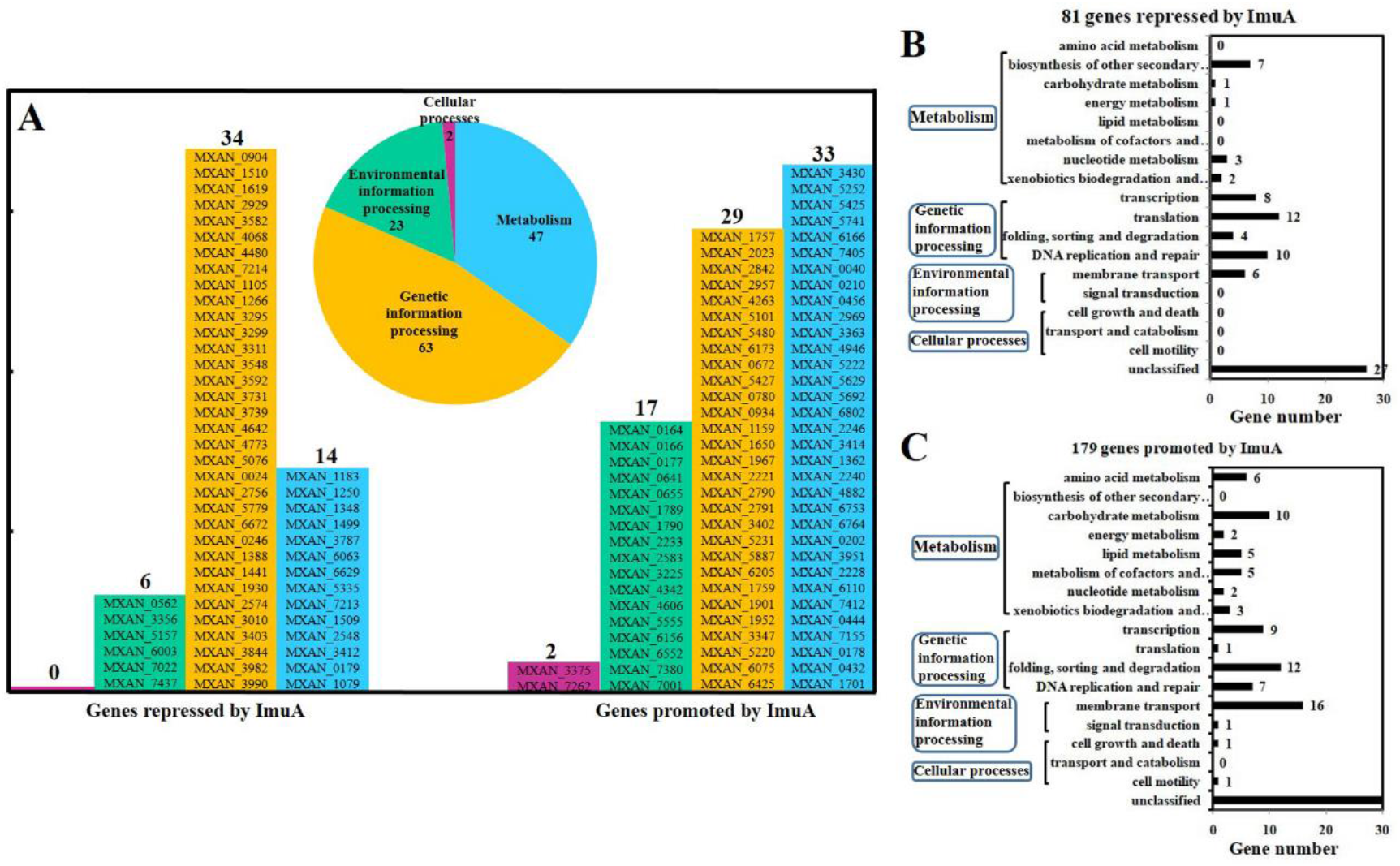
Comparative transcriptome analysis on the *imuA* deletion and overexpression mutants. The transcriptome of wild-type strain was used as the reference and each transcriptome of the three strains was obtained from the average of three repeated sequencing data. The genes differentially expressed in mutants were retrieved. Totally, 260 genes that were synchronously changed in the *imuA* deletion and overexpression mutants were counted. Of the 260 genes, 135 were classified of their functions by KEGG. (A) Classification of 135 function-annotated genes according to KEGG pathway. The genes inhibited and promoted by ImuA are listed in the left and the right panes, respectively. Details of these genes are provided in T**ables S1** and **S2**. (B) GO analysis of the 81 genes inhibited by ImuA (upregulated in the deletion mutant and downregulated in the overexpression mutant). (C) GO analysis of 179 genes promoted by ImuA (downregulated in the deletion mutant and upregulated in the overexpressed mutant).

Furthermore, ImuA-inhibited and ImuA-promoted genes were categorized through the KEGG pathway analysis (**Fig. 4B** and **4C**). The ImuA-inhibited metabolic genes were mainly associated with the biosynthesis of other secondary metabolites (**Fig. 4B**), and ImuA-promoted metabolic genes were in many metabolic processes except the biosynthesis of other secondary metabolites (**Fig. 4C**). Some of ImuA-inhibited genes (**Table S1**) have been proved to be related to resistance in other bacterial species (Desai et al., 2014; Ge & Yu, 2013; Hansen et al., 2008; Kimura et al., 2017; McGeary et al., 2017; Ramirez & Tolmasky, 2017; Tyc et al., 2017; Salinger et al., 2016; Zhang et al., 2002). For instance, *MXAN_5335* was predicted to encode a glycosyl transferase involved in the biosynthesis of polysaccharides, and its homolog in *Aeromonas hydrophila* involved in O-antigen and capsule biosynthesis, which might benefit bacterial resistance to environmental stress (Zhang et al., 2002). The *MXAN_7213* (encoding a peroxidase) and *MXAN_1509* (encoding a diadenosine tetraphosphate hydrolase) are related to oxidative stress and sporulation (Hansen et al., 2008; Kimura et al., 2017).

In genetic information processing, differences were mainly in the pathways of translation, folding-sorting-and-degradation, and replication-and-repair (**Fig. 4B** and **4C**). ImuA showed its inhibitory effect on translation, 12 of the 13 genes involved in translation were inhibited (**Tables S1** and **S2**). In folding sorting and degradation, all 12 ImuA-promoted DEGs were enriched in the protein degradation category and only one of the four ImuA-repressed DEGs were related to protein degradation (**Tables S1** and **S2**). In the replication and repair pathway, 7 DEGs were promoted by ImuA, and10 DEGs were inhibited by ImuA. Notably, *recA1*, *recA2* (Sheng et al. 2020), *dnaE2* and *imuB* were regulated by SOS response and inhibited by ImuA. These outcomes suggested that ImuA was associated with the bacterial SOS response which involved in the induction of secondary metabolites, DNA repair, and mutagenesis, thus resulted in increased bacterial survival rate and adaptation to the altered environment (Baharoglu & Maze, 2014; Kreuzer, 2013; Weissman & Müller, 2010).

### 4. ImuA interacts with ImuB and RecA1

The yeast two-hybrid (Y2H) system was used to identify the ImuA interacting proteins. In the Y2H experiment, ImuA was used as bait to screen its interacting proteins from the proteins near the stalling replication fork, including recombinant proteins (RecA, RecF, RecO and RecR), TLS proteins (ImuB and DnaE2), replication fork binding proteins (DnaE1, SSB and DnaN) and the known SOS response regulator (LexA) from the transcriptomic analysis. The outcome of this analysis showed that ImuA interacted significantly only with ImuB or recombinase RecA1 (**Fig. S3**). Further protein-protein interaction among the three members of ImuA-ImuB-DnaE2 mutasome, and two recombinases (RecA1 and RecA2), proved ImuA could interact with RecA1 and ImuB, not DnaE2. RecA1 interacted with all the other four proteins (ImuA, ImuB, DnaE2, and RecA2), and RecA2 only interacted with RecA1 (**Fig. 5A**). The outcome suggested that RecA1 might be involved in the TLS process.

**Figure 5.**
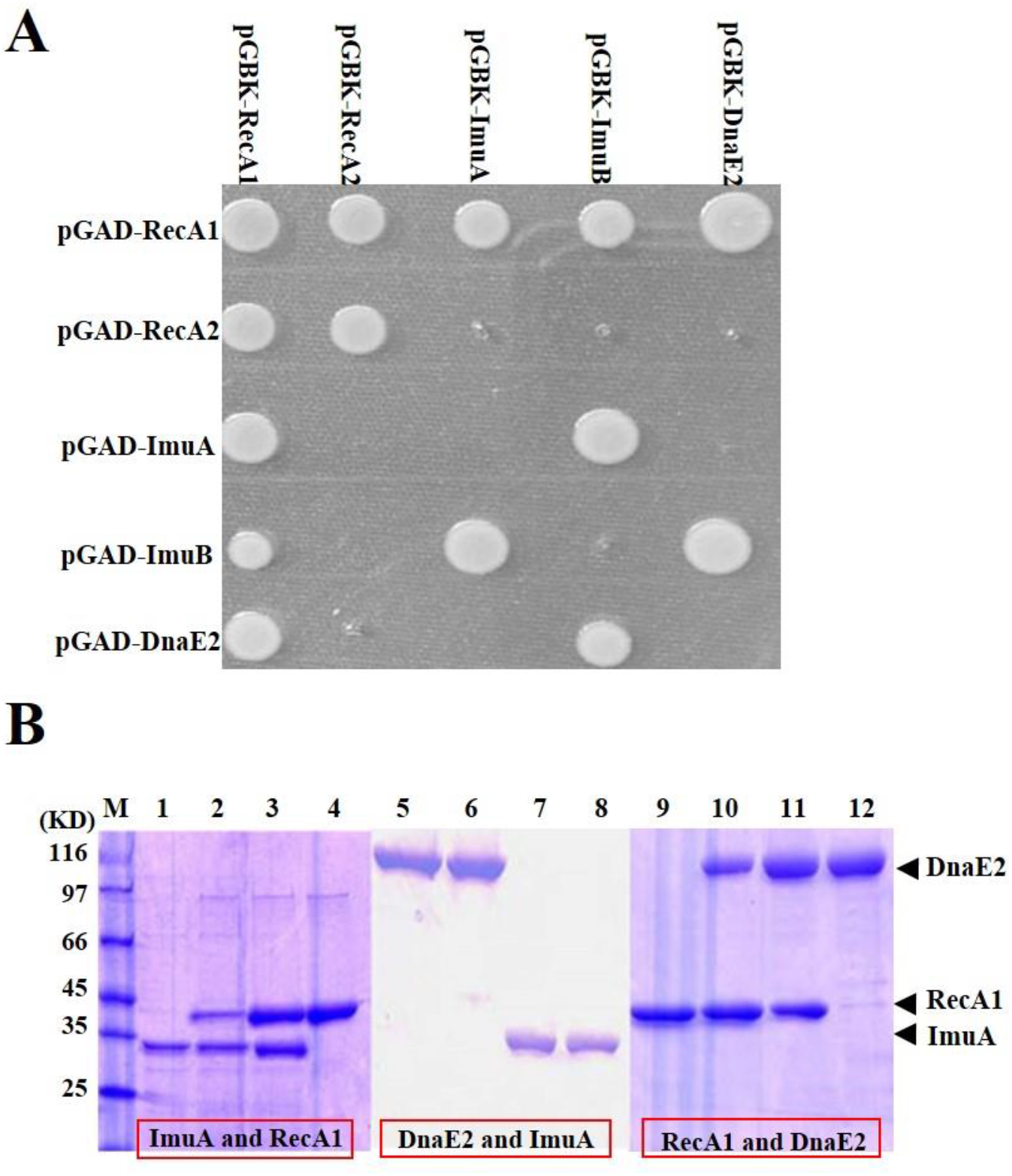
Protein interaction analysis. (A) H2Y analysis on pairwise interactions between RecA1, RecA2, ImuA, ImuB and DnaE2. The genes of these proteins were linked into plasmids pGBKT7 and pGADT7 respectively and the recombination plasmids were pairwise transformed into yeast AH109 to inspect protein interaction on plate with 3-AT. (B) His-Pull down analysis. M, molecular marker; 1, His-ImuA; 2, His-ImuA and RecA1; 3, His-RecA1 and ImuA; 4, His-RecA1; 5, His-DnaE2; 6, His-DnaE2 and ImuA; 7, His-ImuA and DnaE2; 8, His-ImuA; 9, His-RecA1 10, His-RecA1 and DnaE2; 11, His-DnaE2 and RecA1; 12, His-DnaE2.

We further expressed the RecA1, ImuA and DnaE2 proteins with and without a His-tag to verify their interactions using a His-tag pull down assay. As per the SDS-PAGE electrophoresis (**Fig. 5B**), His tagged proteins, *i.e.*, His-ImuA (lanes 1 and 8), His-RecA1 (lanes 4 and 9), and His-DnaE2 (lanes 5 and 12), bound to the nickel column. Besides, RecA1 and ImuA proteins in the combination with His-ImuA (lane 2) and His-RecA1 (lane 3) also bound to the nickel column, respectively. Similarly, RecA1 and DnaE2 proteins interacted to each other (lanes 10 and 11), but ImuA and DnaE2 did not interact (lanes 6 and 7).

### 5. ImuA inhibits the DNA binding and recombinase activities of RecA1

The RecA monomers aggregate on DNA to form nucleoprotein filaments, and containing two polymerization motifs (PM) in the N-terminal and the core ATPase domain (Del et al., 2019; Leite et al., 2016). As an analog of RecA, ImuA has only one conserved PM in the core ATPase domain, and thus could not form RecA-like filaments theoretically (**Fig. 6A** and **S4**), which is consistent with the Y2H result that ImuA could not form self-polymer. Because ImuA has a potential to participate in the RecA filament through the core PM site, thus preventing RecA filament formation and extension (**Fig. S4**).

**Figure 6.**
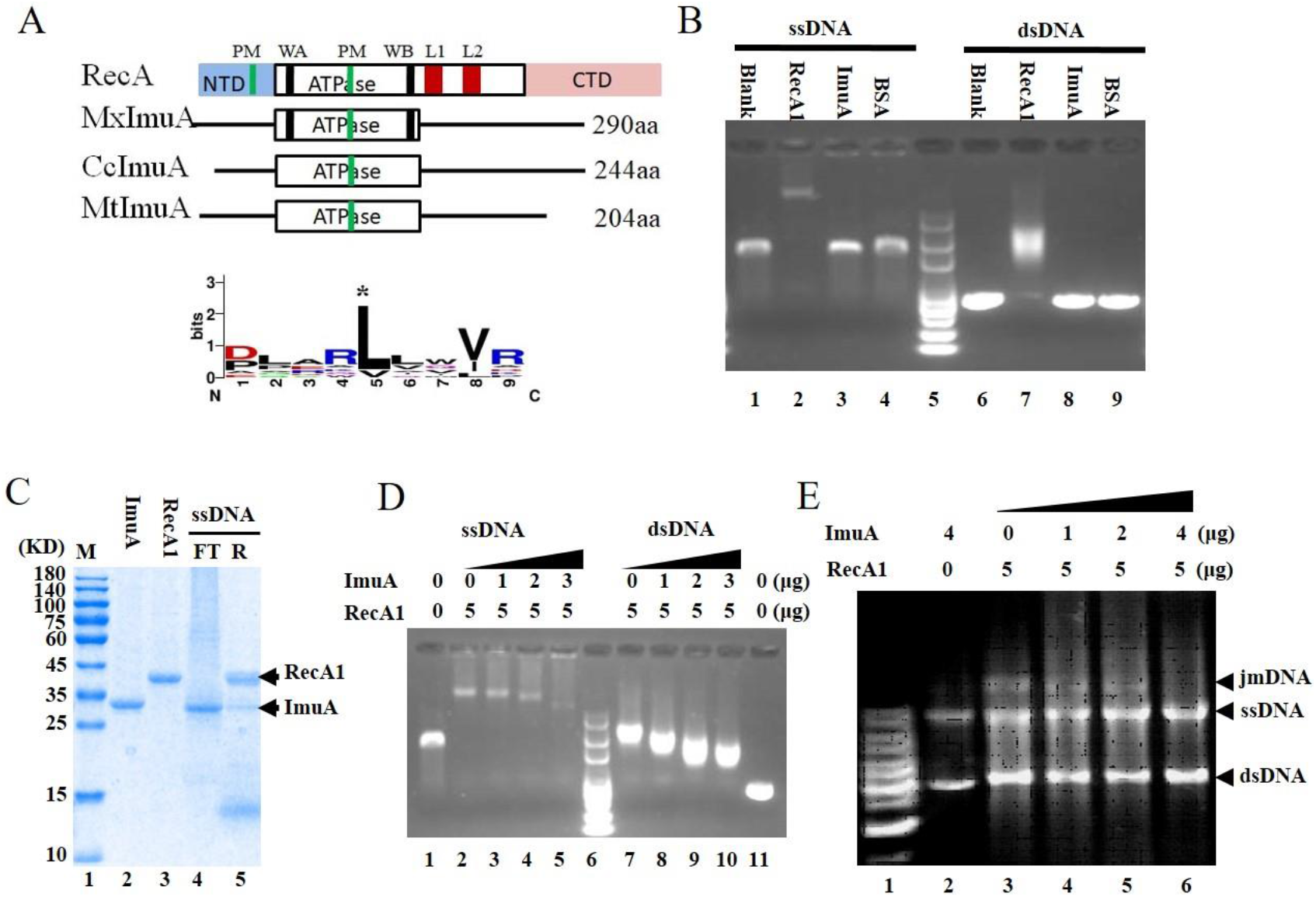
ImuA function analysis. (A) A comparison sketch of ImuAs. RecA, *E. coli* RecA; MxImuA, *M. xanthus* ImuA; CcImuA, *C. crescentus* ImuA; MtImuA, *M. tuberculosis* ImuA; PM, polymerization motif; WA, walker A; WB, walker B; L1 and L2, DNA binding loops 1 and 2. (B) DNA binding ability. RecA1 as positive control and BSA as negative control. (C) ImuA binding ability to the RecA1 coated ssDNA. DNA fixed on Sepharose resin was pre-bound with excess RecA1 protein. ImuA was added to the RecA1 coated ssDNA. SDS page was used to detect the unbound ImuA protein in flow through (FT) and ImuA/RecA proteins bound to resin (R). (D) Effects of increased concentrations of ImuA on the binding of RecA1 to DNA. (E) Effects of ImuA on the recombinase activity of RecA1. jmDNA, joint molecular DNA; ssDNA, single stranded DNA; dsDNA, double stranded DNA.

The ImuA protein lacks the known DNA-binding regions, including the DNA-binding loops and the conserved N-terminus of RecA (**Fig. 6A** and **S4**). We examined the *in vitro* binding ability of ImuA to DNA. Either single-stranded DNA (ssDNA) or double-stranded DNA (dsDNA) incubated with RecA1 lagged significantly, while DNA bands with ImuA or bovine serum protein (negative control) showed no change compared with the blank (no protein added) (**Fig. 6B**). To test whether ImuA could indirectly bind to DNA through RecA1 protein, 12 μg of the ImuA protein (~0.4 nmol) was incubated with 200 ng of ssDNA coated with RecA1 (**Fig. 6C**). The addition of excessive ImuA did not destroy the RecA filament, and the RecA protein was not dissociated clearly (lane 4), but a small amount of ImuA was bound to the resin (lane 5). These findings suggested that ImuA interacted with DNA through RecA1.

A gradient concentration of ImuA proteins was added to the binding reaction solution containing RecA1 and DNA, and the DNA-protein complex was detected using agarose gel electrophoresis (**Fig. 6D**). RecA bound ssDNA or dsDNA to form nucleoprotein filaments (lane 2 or lane 7). The molecular weight and quantity of RecA nucleoprotein filaments decreased (lanes 3-5 or lanes 8-10) with increasing of ImuA concentration. It indicated that ImuA has an inhibition ability on the formation of RecA filaments, probably inhibiting the filament extension by interacting with RecA1.

RecA is a bacterial DNA homologous recombinase that forms a nucleoprotein filament on DNA to catalyze DNA recombination (Del et al., 2019; Leite et al., 2016). RecA1 catalyzed the recombination of circular ssDNA with homologous linear dsDNA, forming a lagging joined DNA band on the electropherogram, while the addition of ImuA to the reaction system significantly reduced this joined molecules (**Fig. 6E**). Thus, ImuA inhibited the RecA-mediated recombination, which was in line with the above DNA binding results that ImuA inhibited RecA binding to ssDNA to form nucleoprotein filaments, and further inhibited DNA recombination.

## Discussion

The sequenced bacterial genomes in the KEGG database were queried for *imuA*, *imuB* and *dnaE2* sequences. A total of 1,669 genomes containing *dnaE2* appeared in the search, of which 1,083 genomes belong to Proteobacteria, 524 to Actinobacteria, and the remaining 62 to Bacteroidetes, Chloroflexi, Deinococcus-Thermus, Firmicutes, and so on. (**Table S3**). Out of the 1,669 *dnaE2*-containing genomes, 1,465 contain an *imuB* gene (87.78% coexistence rate), suggesting the functional correlation between ImuB and DnaE2. Furthermore, out of the 1,465 genomes with *dnaE2* and *imuB*, 1,020 contain a *recA* analog (defined as *imuA*) in front of *imuB*. The coexistence ratio of ImuA, ImuB and DnaE2 is approximately 61.11%, which is highly varied in different taxonomic groups (**Fig. S4**). The presence of ImuA or not significantly affected the SOS mutagenesis catalyzed by DnaE2. For example, in actinomycetes, *Streptomyces coelicolor* lacks *imuA* gene and *S. coelicolor* DnaE2 could not catalyze SOS mutagenesis (Tsai et al., 2012). *M. tuberculosis* encodes ImuA, which plays a key role in DnaE2 catalyzed TLS pathway (Warner et al., 2010).

A total of 1,667 out of 1,669 genomes containing *dnaE2* genes also contained the *recA* gene (**Table S3**). An inevitable competition exists between RecA-catalyzed template switching and DnaE2-catalysed TLS on the stalled replication fork. In *E. coli*, RecA-mediated homologous recombination and Pol V-mediated TLS process compete at the stalled replication fork (Naiman et al., 2016). To prevent homologous recombination, Pol V and RecA formed active mutasome and catalyzed RecA filaments to decompose from the template in the 3’ to 5’ direction in the presence of SSB (Pham et al., 2001). RecA was not directly involved in DnaE2 mutagenesis (Alves et al., 2017), and ImuA undertook the function of inhibiting RecA filaments. So, *M. xanthus* ImuA probably plays two roles in the DnaE2-catalysed TLS, mediating DnaE2 assembly by interacting with ImuB and preventing RecA1-catalysed homologous recombination by interacting with RecA1 (**Fig. 7**). In detail, DNA damage blocks the replication process, exposing ssDNA as a DNA damage signal to activate stalled fork repair. RecA protein was bound to the ssDNA at the stalled replication fork to form nucleoprotein filaments to initiate template switching pathway. Binding of ImuA to the RecA filament hindered the extension of filament and the recombination ability of RecA, thus inhibiting template switching. Meanwhile, the interaction between ImuA and ImuB facilitated the assembly of ImuB-DnaE2 dimer onto the DNA replication fork, and initiated TLS pathway.

**Figure 7.**
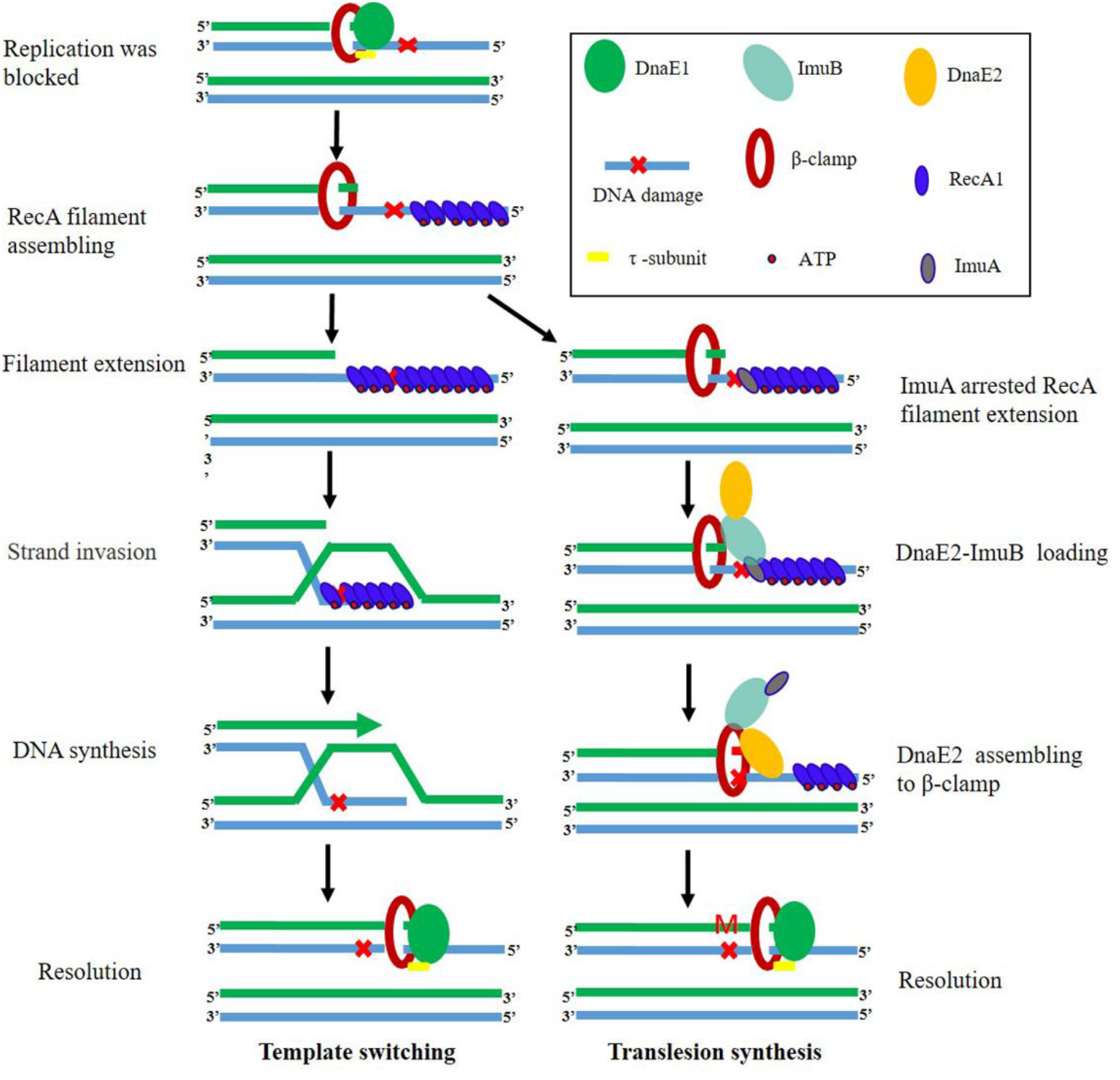
Model of ImuA function in template switching and TLS in *M. xanthus*. Bacteria have two pathways to rescue stalled replication forks, error-prone TLS and error-free template switching. When replication encounters a DNA damage and is blocked, ssDNA is exposed as a DNA damage signal to activate SOS response. The SOS genes, including *recA1*, *imuA*, *imuB*, and *dnaE2*, are induced. RecA protein binds to the ssDNA at the stalled replication fork to form nucleoprotein filaments and initiates template switching pathway. At the same time, the DnaE2 approaches the stalled replication fork to initiate the TLS pathway. Combination of ImuA and RecA hinders the extension of RecA nucleoprotein filaments and the recombinase activity of RecA, thus inhibiting template switching. ImuB guides DnaE2 to assemble on the replication fork by binding to ImuA/β-clamp to start TLS. In the absence of ImuA, RecA binds to the exposed ssDNA to form a filament, which promotes the error-free template switching.

*M. xanthus* RecA1 is a classical bacterial RecA involved in DNA recombination and LexA-dependent SOS induction. ImuA interacted with RecA1 to prevent the extension of RecA1 filaments, which might explain the effects of ImuA on UV sensitivity and SOS gene expression (**Fig. 3 and 4**). As the intracellular level of RecA1 is very low (Norioka et al., 1995; Sheng et al., 2020). Thus, the RecA1-catalysed template switching accounts for a relatively small proportion in the recovery of the stalled DNA replication process, and the effect of inhibiting RecA1 to improve the DnaE2-catalysed mutagenesis is also insignificant, which may be a reason why the deletion of *M. xanthus* ImuA had no obvious effect on the bacterial mutation frequency. Another plausible reason might be that ImuA deletion downregulated the mutagenesis activity of the DnaE2 protein and simultaneously upregulated the transcription of *dnaE2* through SOS response.

ImuA also interacts with ImuB; which is required for the DnaE2 assembly (Warner et al., 2010). ImuB is a homolog of the Y family protein UmuC and has the potential to bind to RecA/ImuA/β-clamp (Del et al., 2019; Timinskas & Venclovas, 2019). In the absence of ImuA, DnaE2 assembly on the DNA might bemediated by ImuB and RecA1 or β-clamp interaction. The binding of ImuB with RecA/ImuA was probably through the C-terminal tail of UmuC (similar RecA-NT motif) and the core ATPase domain (core PM site) of RecA/ImuA similar to that of the RecA filament (Timinskas & Venclovas, 2019). Thus, ImuB may detach the RecA/ImuA protein from the filaments in the 3’-5’ direction by competitive binding to the core PM site of RecA/ImuA (**Fig. 7**); which demands an in-depth investigation.

## Materials and Methods

### Bacterial strains and growth conditions

Details of bacterial strains, plasmids, and oligonucleotides used in this study are provided in **Table S4** and **S5**. The *M. xanthus* strains were cultivated in Casitone-based CYE medium for growth assays and on TPM agar for developmental assays. *E. coli* strains were routinely incubated on Luria-Bertani (LB) agar or in LB liquid broth. *M. xanthus* strains were incubated at 30 °C, and *E. coli* strains were incubated at 37 °C. Solid or liquid medium was supplemented with 33 μg/ml of kanamycin (Kan) or 100 μg/ml of ampicillin (Amp) as required.

### Gene mutation and overexpression of *M. xanthus imuA*

Markerless deletion mutation was carried out to knockout *imuA* or *imuB* gene in *M. xanthus* DK1622 using pBJ113 plasmid. The plasmid contained a kanamycin resistant cassette for the first round of screening and a *galK* gene for the negative screening (Julien et al., 2000). To avoid the potential effect on the downstream gene expression, the middle sequence of *imuA* gene was deleted and 9 bp from N-terminal and 93 bp from C-terminal were retained (**Fig. S2**). Similarly, the middle sequence of *imuB* was deleted and 54 bp of N-terminal and 99 bp of C-terminal was kept (**Fig. S2**). Briefly, up- and down-stream homologous arms were amplified using primers (**Table S5**) and ligated using the *Bam*H1 site. The fragment was inserted into the *Eco*RI/*Hin*dIII site of pBJ113. The resulting plasmid was introduced into *M. xanthus* strains via electroporation method (1.25 kV, 200 W, 25 mF, 1 mm cuvette gap). The second round of screening was performed on CYE plates containing 1% galactose (Sigma). The *imuA* mutant (named IA) and *imuB* mutant (named IB) were identified and verified by PCR amplification and sequencing (**Fig. S2**).

To overexpress *imuA* gene, the whole *imuA* gene, along with its promoter, was amplified and cloned into the *M. xanthus* integrative vector pSWU19 (Wu & Kaiser, 1997). The recombinant plasmid was transformed into *M. xanthus* DK1622 cells. Individual kanamycin-resistant clones were selected and validated using PCR amplification. We also transformed the empty vector into *M. xanthus* wild type strain as a control.

Restriction enzymes, DNA ligase, and other enzymes were used according to the manufacturers’ instruction. All DNA products were validated using DNA sequencing.

### Growth and mutation frequency assays

*M. xanthus* cells in exponential phase were used to determine cell growth curves. For UV irradiation, 15 ml of cell suspension in 130 mm glass Petri dishes was stirred gently with a magnetic rod and irradiated with UV rays (254 nm) at room temperature. Then, the cells were diluted in fresh medium and incubated at 30 °C. The density of the cells was measured every 12 h to generate the growth curves.

The mutation frequency assay was conducted by screening the nalidixic acid resistant strains, as described previously (Tzeng et al., 2006). Briefly, cells were mock-irradiated or irradiated with 15 J/m^2^ UV. After a 4-h acclimation period, approximately 5 × 10^9^ *M. xanthus* cells were placed on CYE agar containing 40 μg/ml nalidixic acid incubated for 5 days. Nalidixic acid-resistant candidates were counted and mutation frequency was calculated using the following formula, mutation frequency = nalidixic acid resistant clones / 5 × 10^9^.

### RNA extraction, RT-PCR and RNA sequencing

*M. xanthus* cells were UV irradiated at the dose of 0, 5, and 15 J/m^2^, and then, these cells were diluted in fresh medium and incubated for 0-7 h. Total RNA from these cells was extracted using RNAiso Plus reagent (Takara, Beijing) following the manufacturer’s protocol. RNA concentration was quantified using Nanodrop 2000 (Thermo Fisher Scientific, USA). The cDNA was synthesized using PrimeScript RT Reagent Kit (Takara, Beijing). Genomic DNA (gDNA) in the RNA sample was removed with gDNA Eraser (Takara, Beijing). The first-strand cDNA was synthesized by PrimeScript RT Enzyme following the manufacturer’s instructions with random primers.

The resulting cDNA was diluted 5 X before performing RT-PCR. The primers designed for *imuA*, *dnaE2*, *recA*1 and *recA*2 genes are listed in **Table S5**. RT-PCR was accomplished using the SYBR premix Ex Taq kit (Takara, China) on an ABI StepOnePlus Real-Time PCR System (Thermofisher Scientific, USA) with the following program: 3 min at 95 °C, followed by 40 cycles of 30 s at 95 °C, 30 s at 55 ° C, and 15 s at 72 °C. The relative quantification of mRNAs of interest was performed by the comparative CT (2^−ΔΔCT^) method, using the *gapA* (glyceraldehyde-3-phosphate dehydrogenase) gene as an endogenous control, as described previously (Peng et al., 2017). The mock-treated samples (0 J/m^2^) were used as control.

RNA sequencing was conducted by Vazyme. Purified double-stranded cDNA were end-repair, added bases A, and ligated with sequencing adaptors. After PCR amplification, the cDNA was sequenced on an Illumina HiSeq 4000. All the upregulated and downregulated genes were got by comparing with control, and their gene functions were explored using database annotations such as GO, and KEGG.

### Expression, purification and pull-down analysis of ImuA, DnaE2 and RecA1

Genes *imuA* and *imuB* were cloned into *E. coli* expression vectors pET15b and pET3a, respectively. Then these recombinant plasmids were transformed into *E. coli* BL21(DE3) and cultured in LB medium at 37 °C. Induction of expression was initiated at an OD_600_ = 0.6 by the addition of IPTG (1 mM). Cells were harvested 3 h later and sonicated to prepare the crude extract. 6x Histagged proteins were purified from the crude extract using Ni-NTA agarose columns (Qiagen) according manufacturer’s instructions, and no 6xHis tagged proteins were purified by salting out and ion exchange chromatography.

In the His-pull down analysis, purified His-tagged proteins were first incubated with Ni-NTA agarose beads. Subsequently, the beads were extensively washed and incubated with no His-tag labeled protein. The proteins bound to the beads were eluted with 150 mM imidazole and separated by SDS-PAGE.

### Electrophoretic mobility shift assay

To test DNA binding ability, the electrophoretic mobility shift assays were performed in the reactions containing 25 mM Tris-HCl, pH 7.0, 50 mM NaCl, 4% glycerol, 1 mM DTT, 10 mM MgCl_2_, 1.5 mM ATP, M13 circle ssDNA or 3 kb dsDNA (derived from M13) and protein (DNA and protein concentrations as depicted in the figures below). The solution was incubated at 32 °C for 20 min, and electrophored on 1% agarose gel for 2 h, at 2 V cm^−1^ and 4 °C. This gel was stained with Ethidium Bromide. This experiment was for three times, and a representative blot is shown below.

### DNA strand exchange reactions

The RecA-dependent DNA strand exchange reaction was carried out as described (Shan et al., 1996) between M13 circular ssDNA and the linear dsDNA (3 kbp fragment derived from M13). The reactions were carried out at 30 °C in SEB solutions containing 25 mM Tris-HCl, pH 7.0, 50 mM NaCl, 4% glycerol, 1 mM DTT, 10 mM MgCl_2_, and an ATP-regenerating system (10 units/ml of pyruvate kinase and 3.3 mM phosphoenolpyruvate). After preincubation of ssDNA with RecA1 protein at 30 °C for 5 min, the reaction mixture was supplemented with 1.5 mM ATP and ImuA, and continued 5-min incubation. Linear dsDNA was added to start the DNA strand exchange reactions. The reactions were stopped by the addition of 5 μl of gel loading buffer (0.125% bromophenol blue, 25 mM EDTA, 25% glycerol, and 5% SDS) to reaction mixture. Furthermore, these samples were electrophoresed in 0.8% agarose gel with TAE buffer.

### Yeast two hybrid

The yeast two hybrid phenotypic assay to evaluate protein interaction was performed as described previously (Fields & Sternglanz, 1994). Genes *imuA*, *imuB*, *dnaE2*, *dnaE1*, *recA1*, *recA2*, *recO*, *recR*, *dnaN*, *lexA* and *ssb* were cloned into pGBKT7 and pGADT7 plasmids. These plasmids were transformed into yeast AH109 cells and interacting phenotypes were screened for their ability to grow on the selective medium with 3-AT. These experiments were repeated three times.

### Binding analysis of ImuA to RecA coated ssDNA

M13mp18 ssDNA was immobilized on CNBr-activated Sepharose 4B (GE), as described previously (Kingston et al., 1999). 20 μl of the agarose beads containing 200 ng M13 (+) strand (~0.6 nmol) were mixed with 14.4 μg RecA1 protein (~0.4 nmol) in 200 μl of SEB buffer for 5 min at 30 °C by constant tapping. Theoretically, one RecA molecule binds to three nucleotides, and 14.4 μg RecA1 is twice the amount of RecA required to theoretically bind 200 ng of DNA. The beads were captured by centrifugation at low speed and washed twice with SEB buffer to remove unbound proteins. Then, 12 μg ImuA protein was loaded to the RecA-DNA-beads and continue incubation at 30 °C for 5 min. The unbound protein was washed out with SEB buffer. The DNA binding proteins on beads were cut using the ultrasonic treatment, collected with centrifugation and separated by SDS-PAGE.

## Acknowledgements

This work was financially supported by the National Natural Science Foundation of China (NSFC) (Nos. 31670076 and 31471183), National Key research and development Programs of China (Nos. 2018YFA0900400 and 2018YFA0901704), the Key Program of Shandong Natural Science Foundation (No. ZR2016QZ002), Special investigation on scientific and technological basic resources (No. 2017FY100302) to YZL and Key Research & Developmental Program of Shandong Province (No. 2019JZZY020308) to DHS.

## Data availability statement

All datasets generated for this study are included in the manuscript and the supplementary files.

## Supplementary figures

**Figure. S1.**
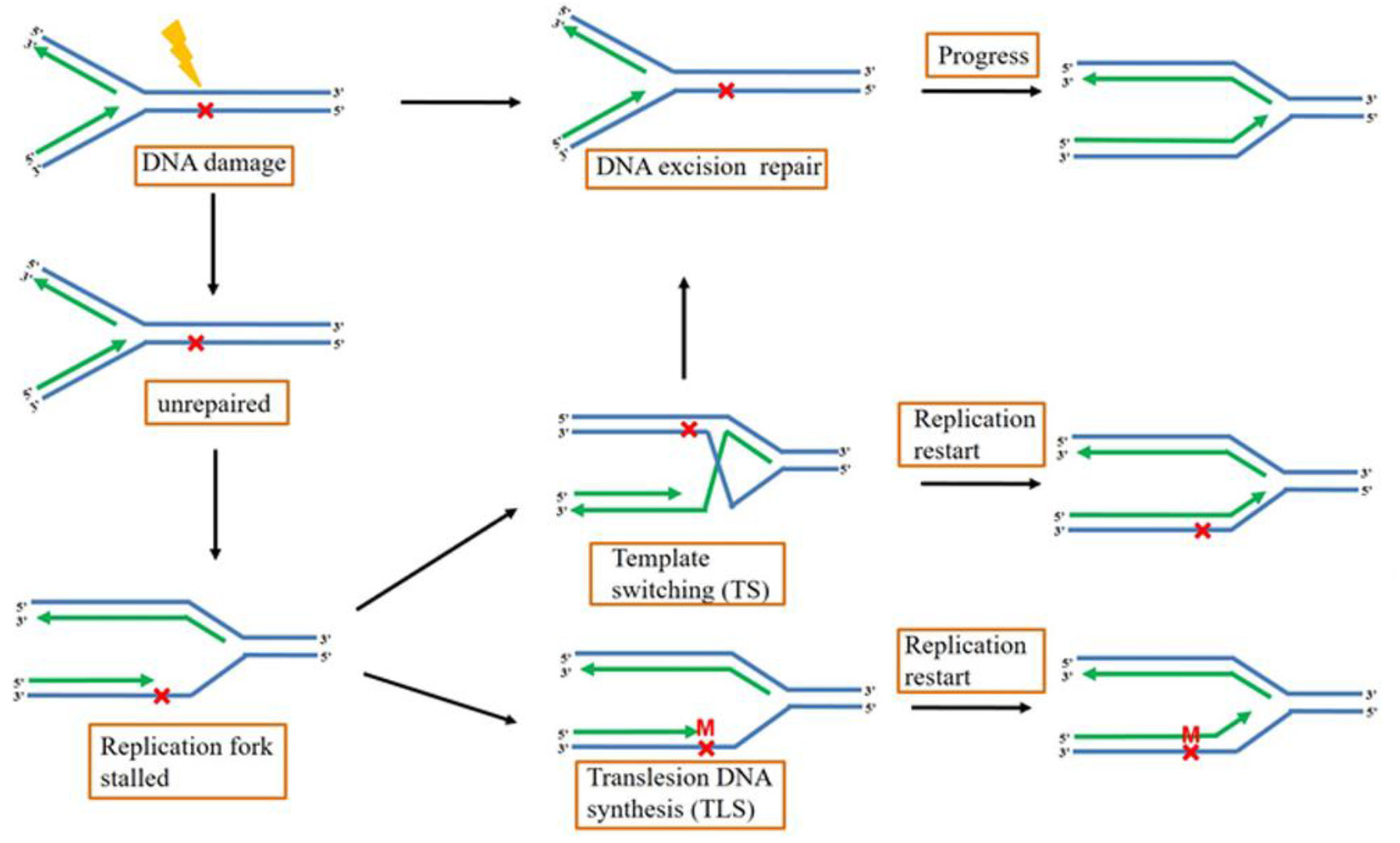
Schematic diagram illustrating two bacterial pathways for restarting replication forks. The majority of base-specific damage is repaired by DNA repair before replication. DNA lesions that is not repaired in time or near the replication fork will stall replication. Two pathways for stalled fork recovery, error-prone translesion DNA synthesis (TLS) and error-free template switching (TS), can bypass the lesions in order to restart DNA replication downstream. Newly synthesized DNA strand and their templates are shown in green and blue, and 3′-ends of growing DNA chains are shown by arrows. The sites of DNA damage (×) and mutation (M) were shown in red.

**Figure S2.**
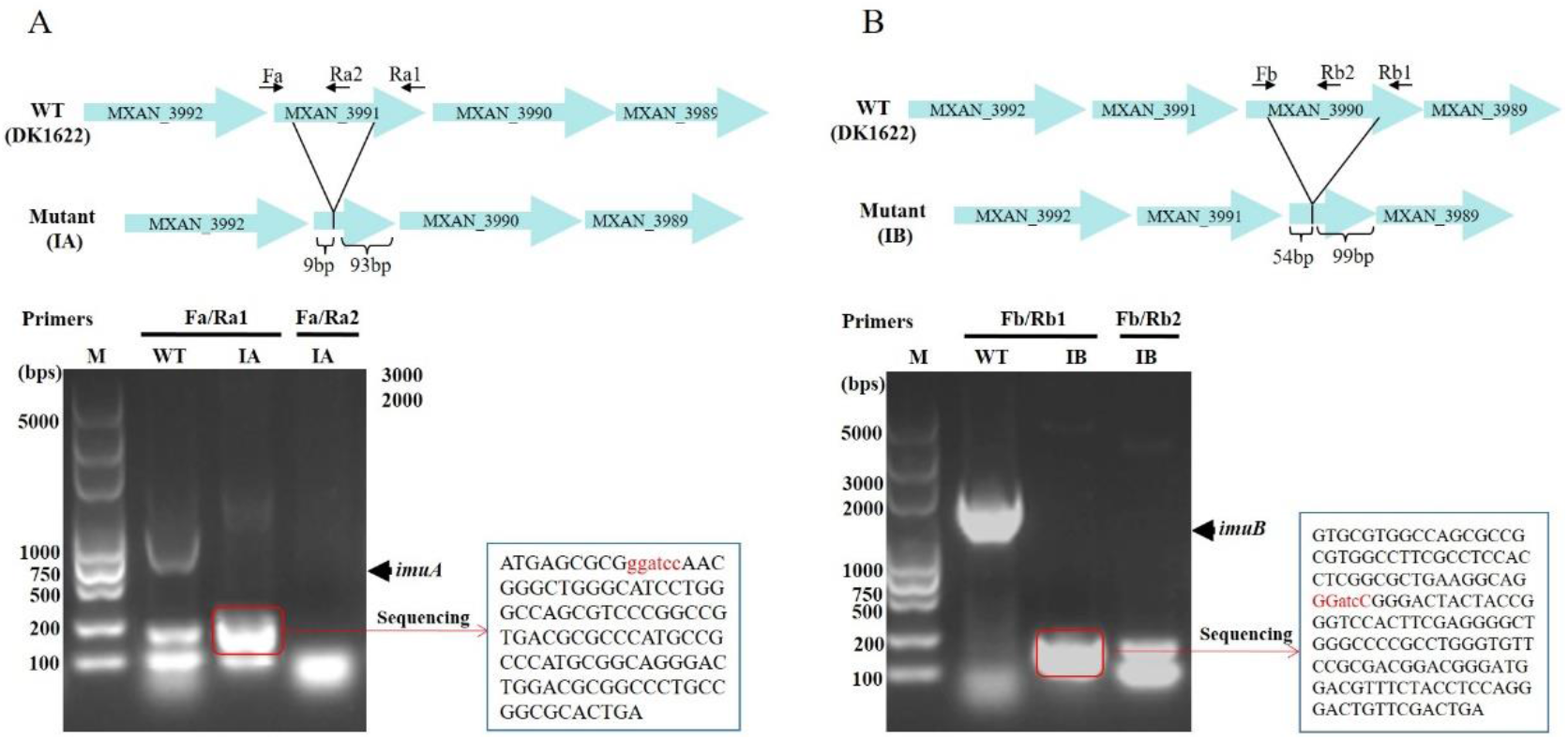
Marker-free gene knockout *of imuA* (A) and *imuB* (B) in *M. xanthus* DK1622. The schematic maps of the location for gene knockout are provided as the upper panel. In the process of gene knockout, the N- and C-terminal respectively retain some bases linked by a *Bam*H1 recognition sequence (ggatcc). The lower panel is the PCR validation of the mutants with primers indicated by black arrows in upper schematic maps. Further sequencing results of the PCR product from the mutants are shown aside and the N-and C-terminal residues are linked by *Bam*HI sites (red)

**Figure S3.**
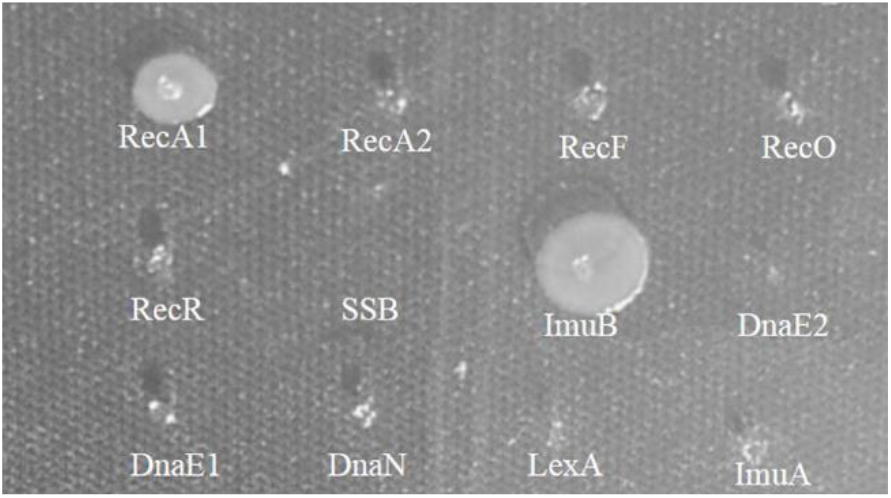
Using ImuA as a baiting protein to screen its interacting proteins by Y2H method. ImuA was cloned into pGBKT7 plasmid and used as a bait to screen its interacting proteins among the known replication and recombination proteins in pGADT7 in AH109.

**Figure S4.**
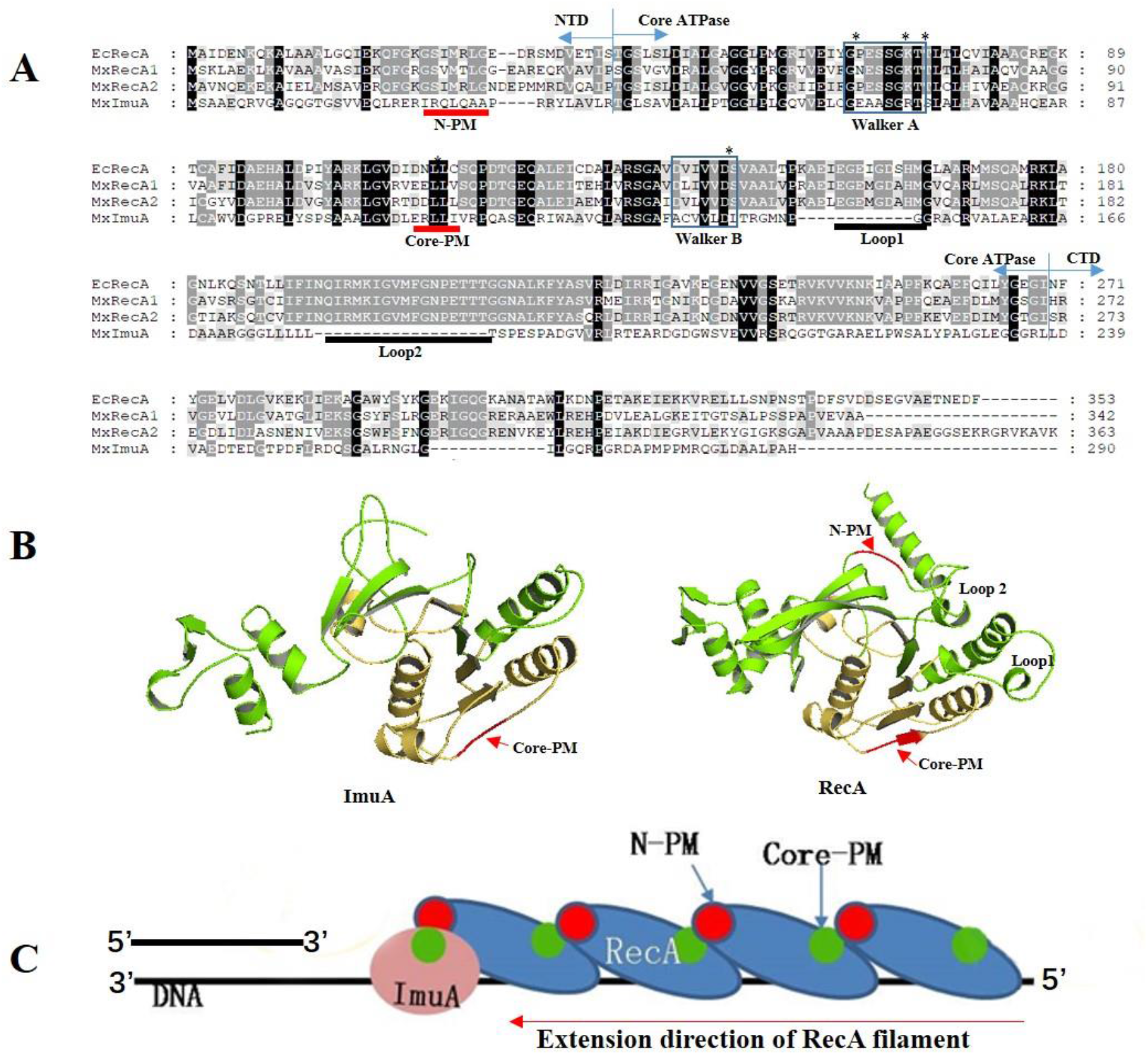
Sequence and structure analysis of *M. xanthus* ImuA. (A) Sequence alignment of *M. xanthus* ImuA (MxImuA) with RecA1 (MxRecA1), RecA2 (MxRecA2) and *E. coli* RecA (EcRecA). The ATP binding Walker A and B motifs are marked in blue frame, the putative DNA binding sites (Loop 1 and Loop 2) are underlined in black and the polymerization motifs (N-PM and Core-PM) are underlined in red. Conservative amino acids of Walker A, core-PM and Walker B motifs were marked with asterisks. (B) Three dimensional structure model of *M. xanthus* ImuA, compared with RecA1. (C) Predicted interaction diagram of *M. xanthus* ImuA with RecA1. N-PM and core-PM are the binding sites located at N-terminal and core ATPase domain of RecA to form polymers.

**Figure S5.**
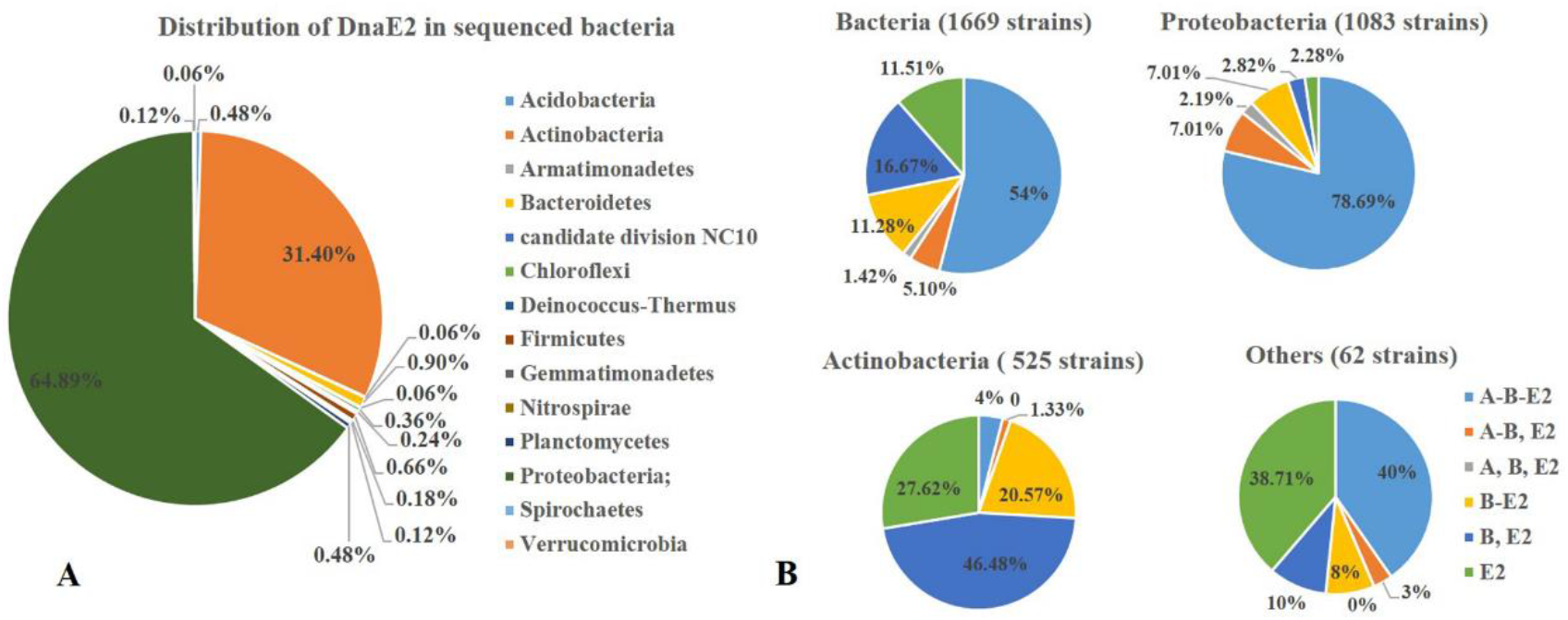
Distribution and location relationships of *imuA*, *imuB* and *dnaE2* genes in sequenced bacterial genome. (A) Distribution of DnaE2. DnaE2 is distributed in 14 bacterial phyla. (B) Location relationship of *imuA, imuB* and *dnaE2* in the 1669 genomes carrying *dnaE2*. A-B-E2: *imuA*, *imuB* and *dnaE2* genes coexist in the same genome and are adjacent to each other; A-B,E2: *imuA*, *imuB* and *dnaE2* genes coexist in the same genome and *imuA*-*imuB* are adjacent to each other, *dnaE2* is separated; A,B,E2: *imuA*, *imuB* and *dnaE2* genes coexist in the same genome and are separated to each other; B-E2: *imuB* and *dnaE2* genes coexist in the same genome and are adjacent to each other, *imuA* disappears; B,E2: *imuB* and *dnaE2* genes coexist in the same genome and are separated to each other, *imuA* disappears; E2: only a *dnaE2* gene, *imuA* and *imuB* disappear.

## Supplementary tables

**Table S1.**
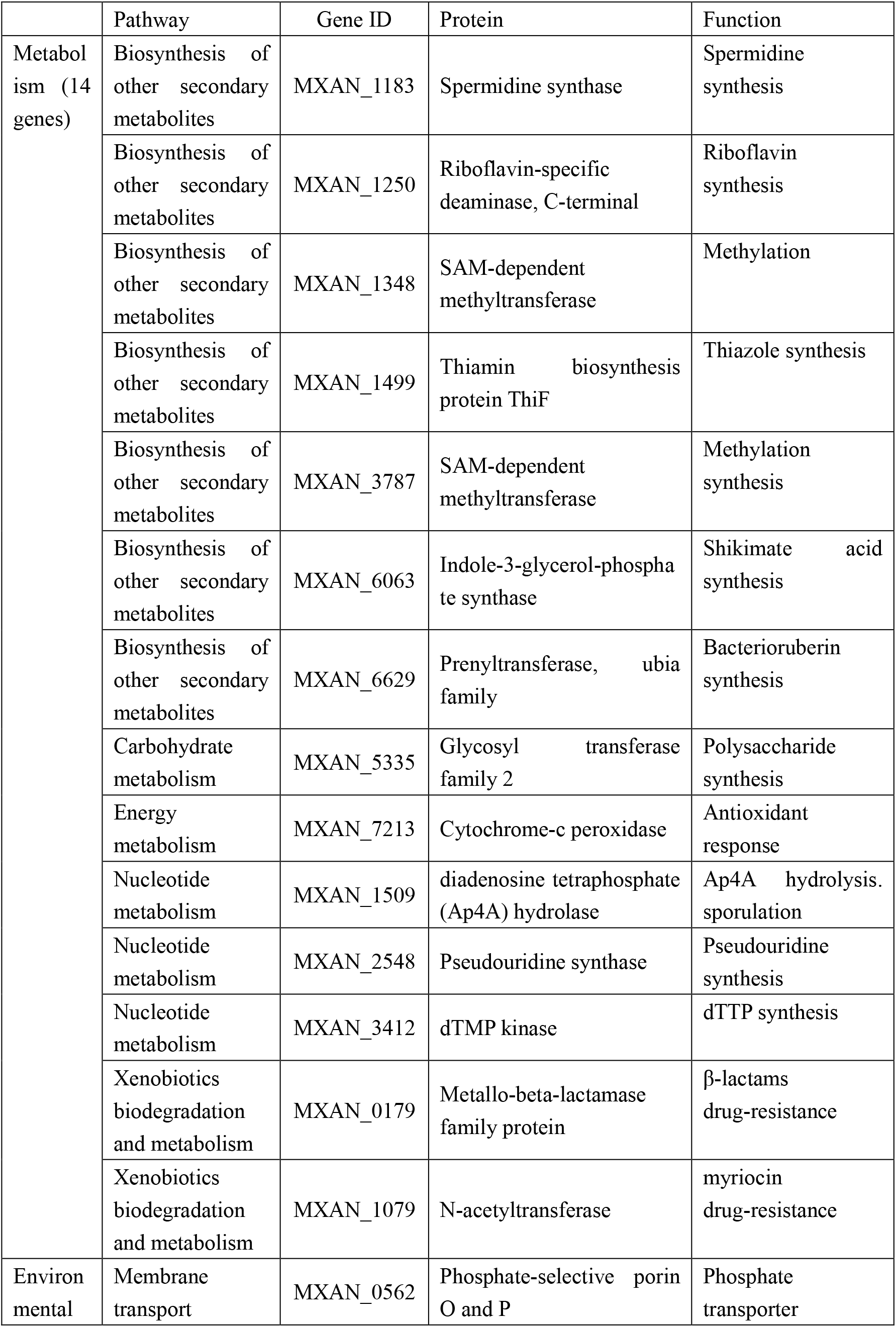
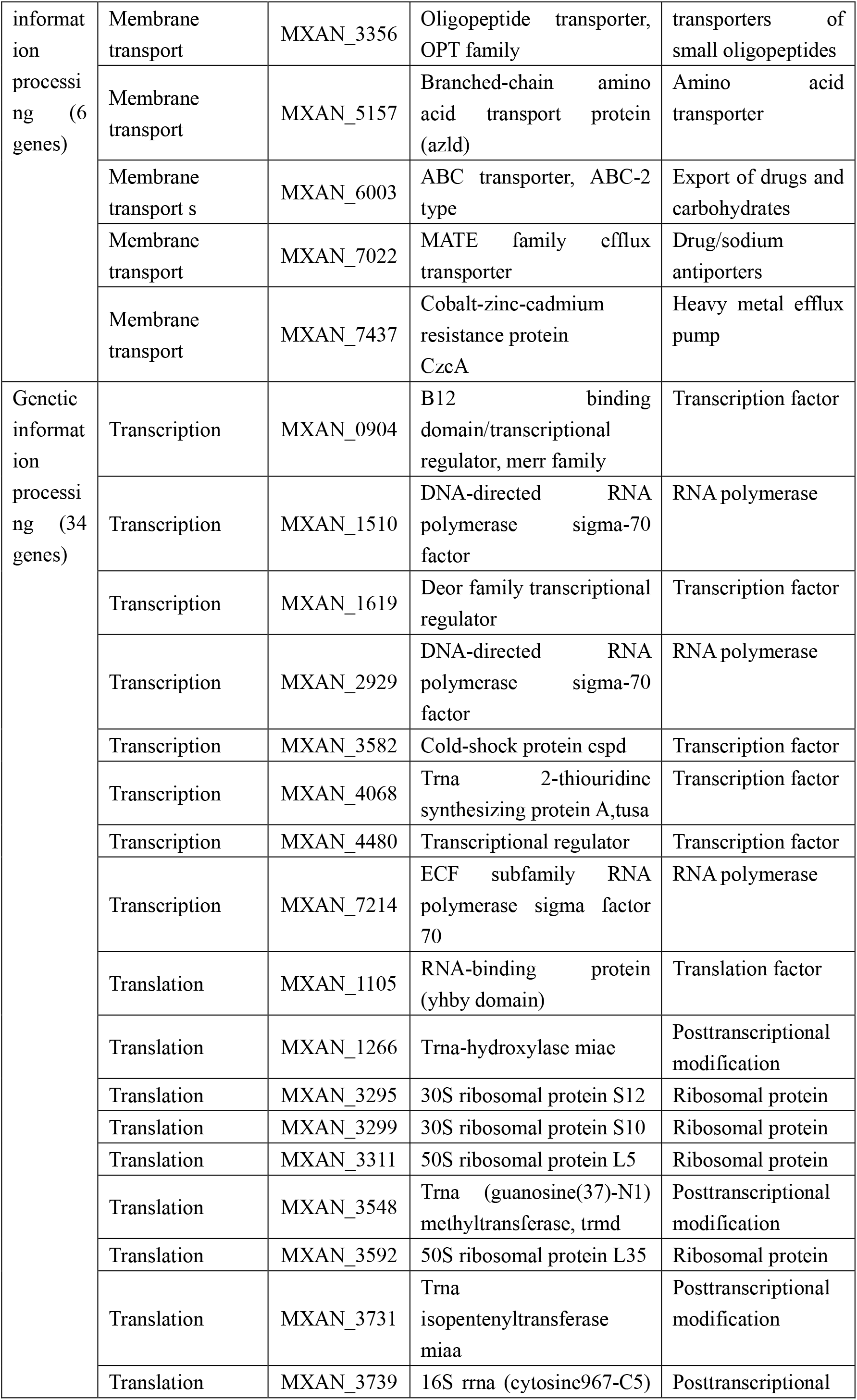
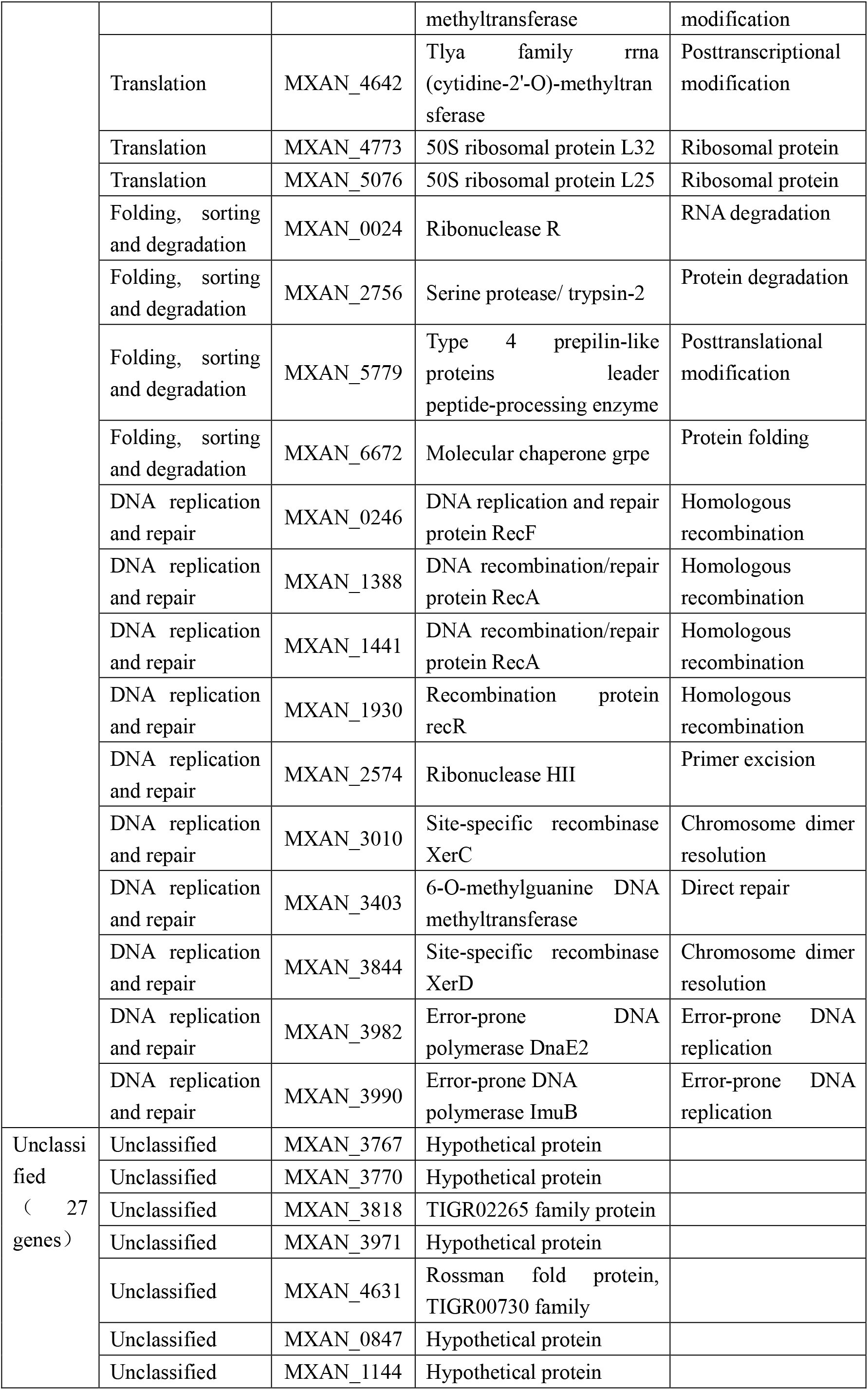
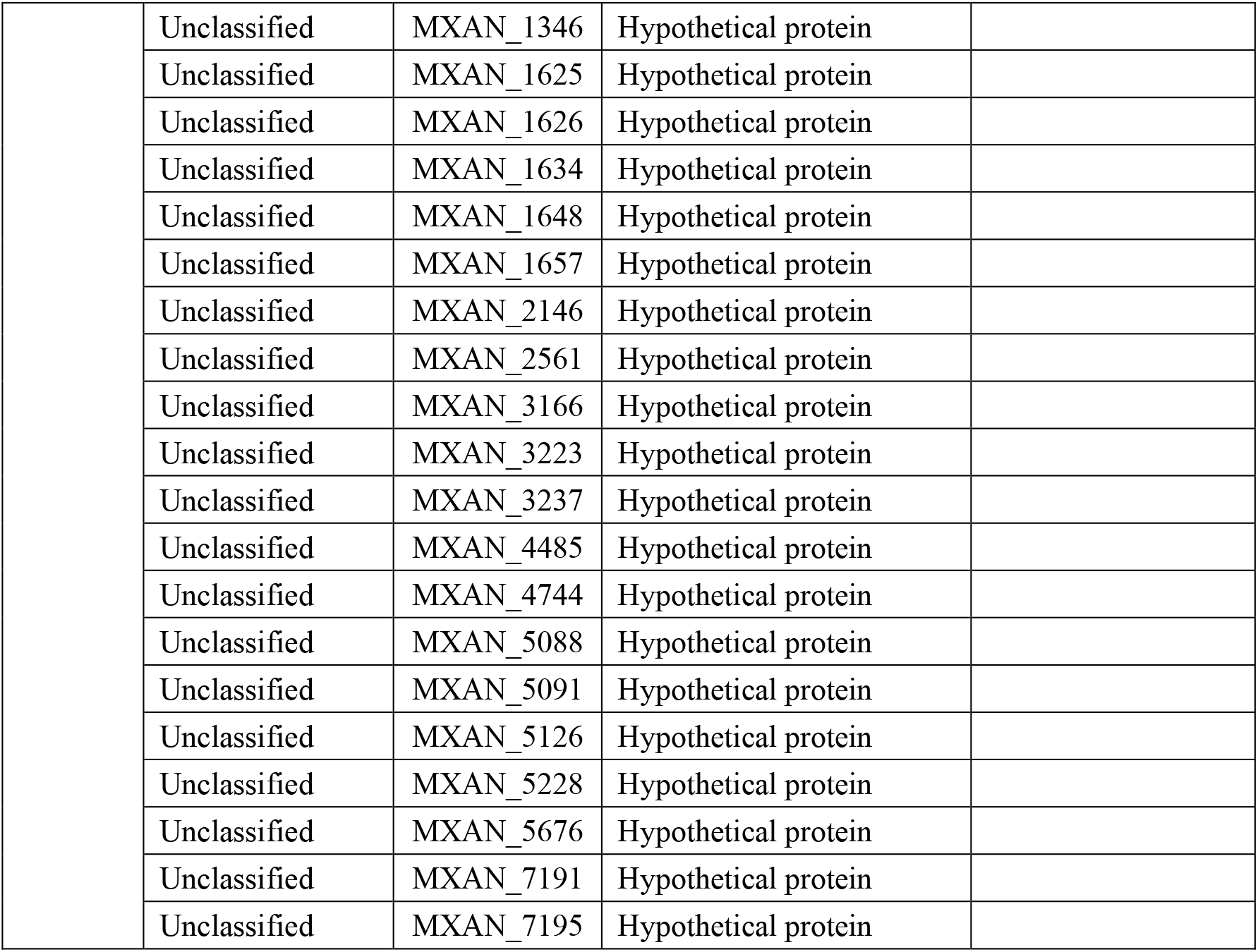
81 genes inhibited by ImuA

**Table S2.**
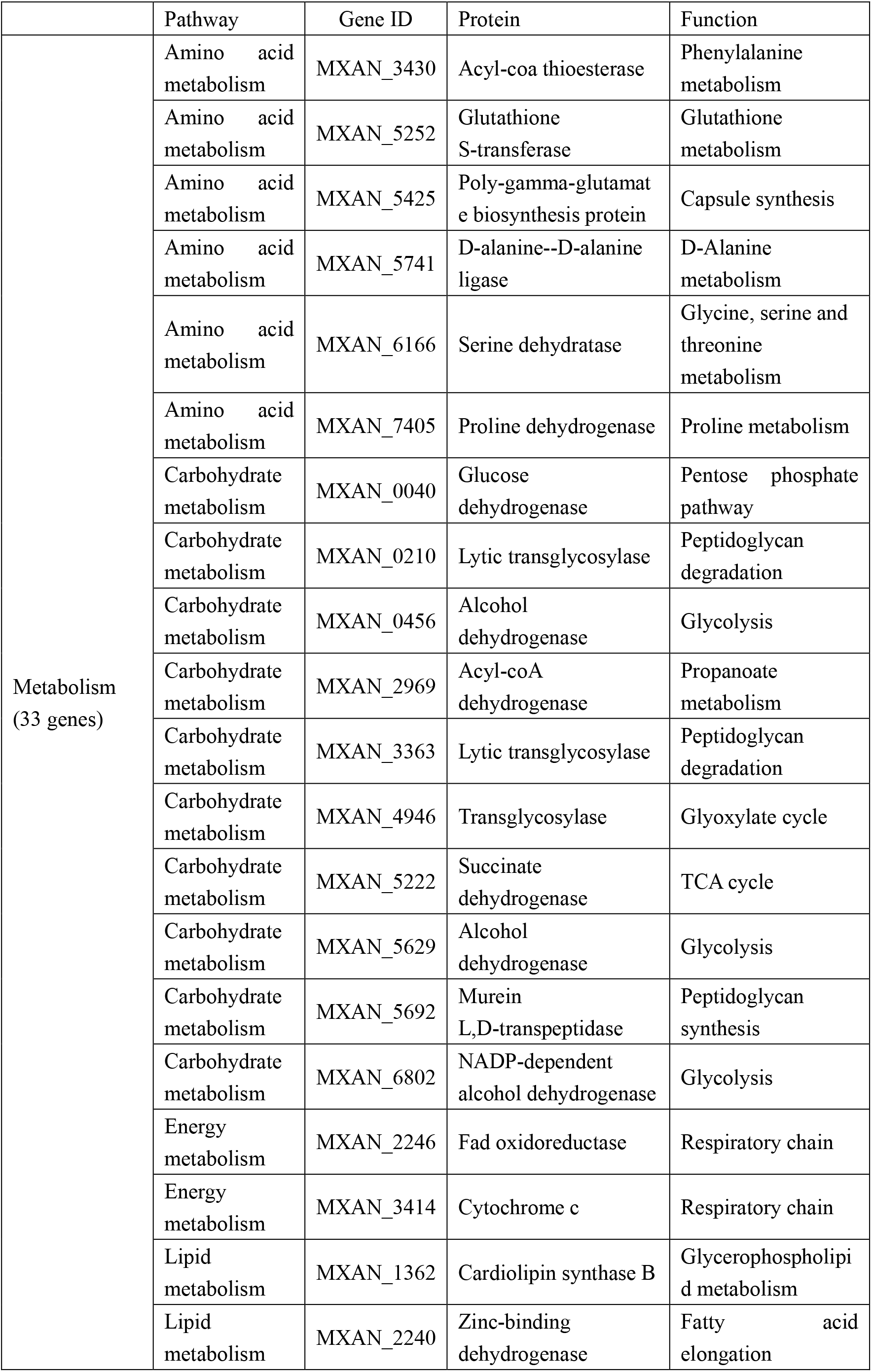
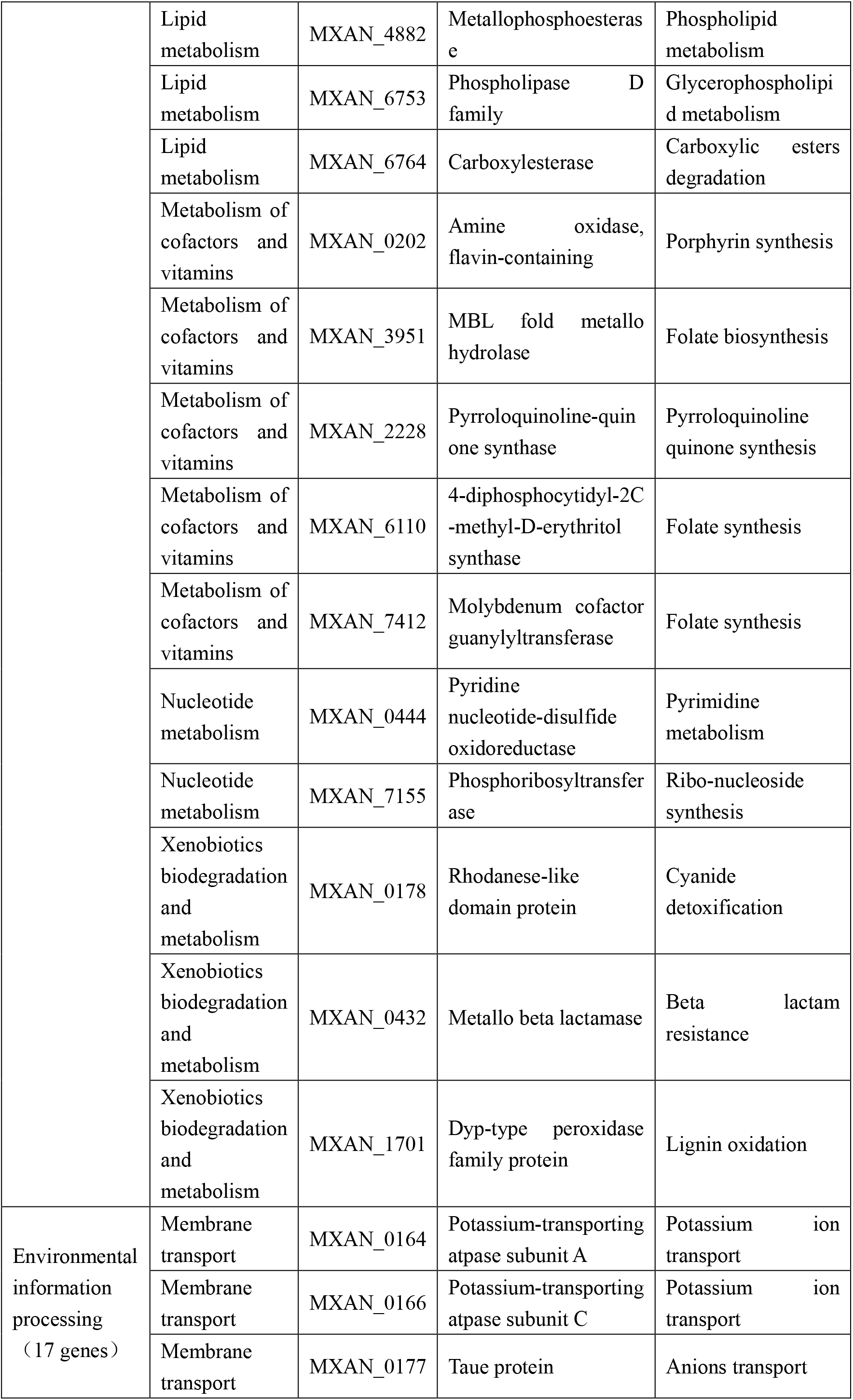
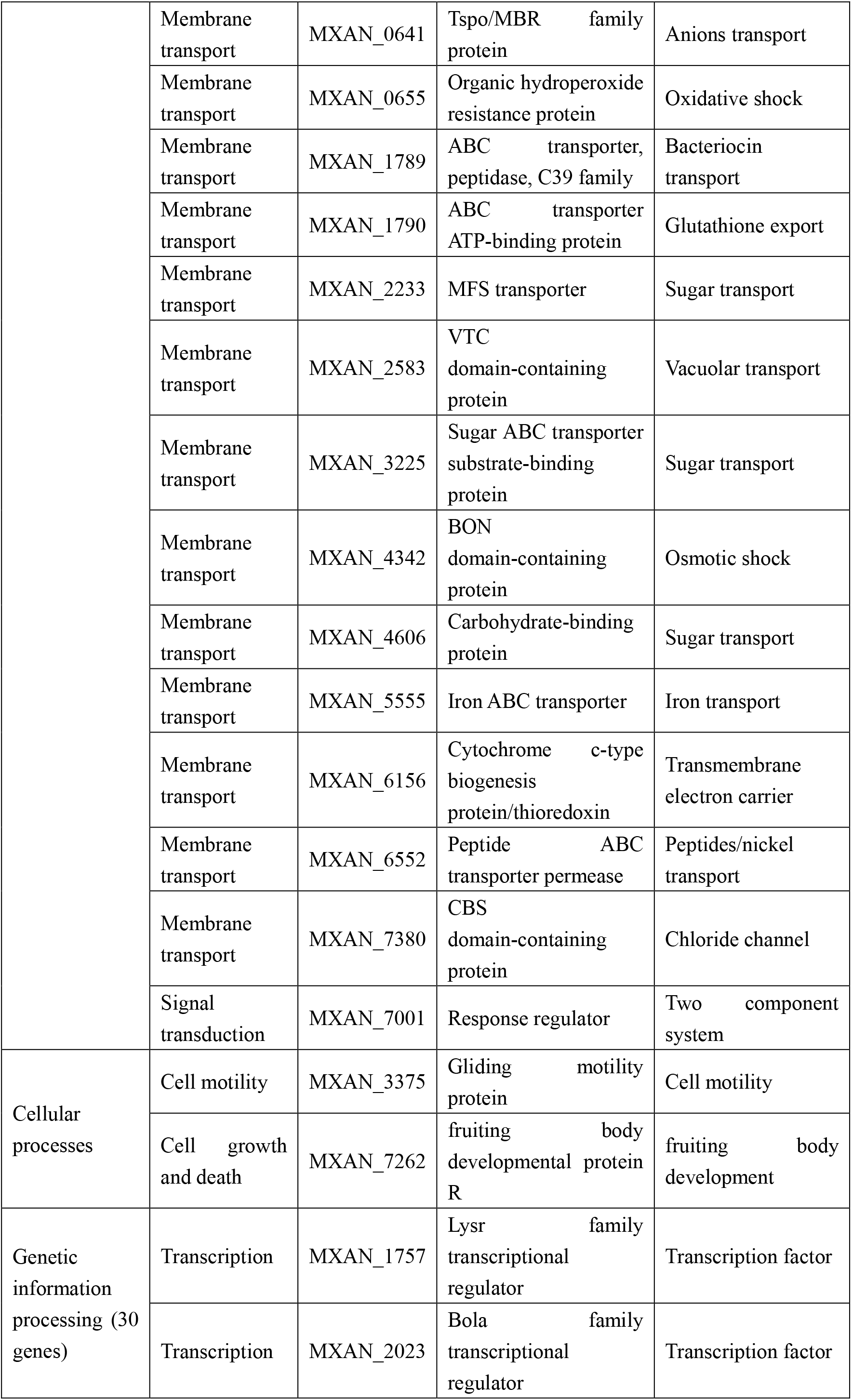
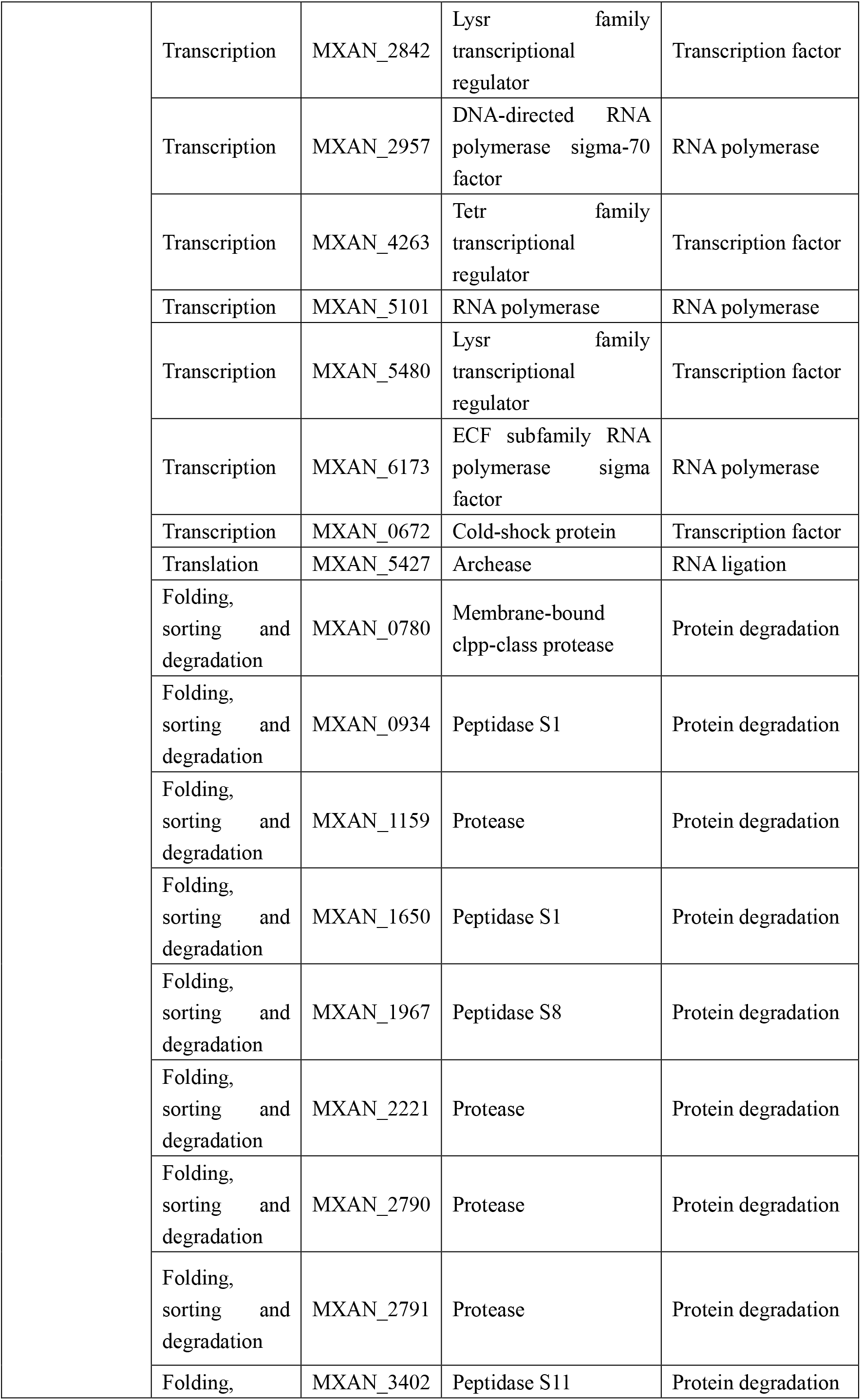
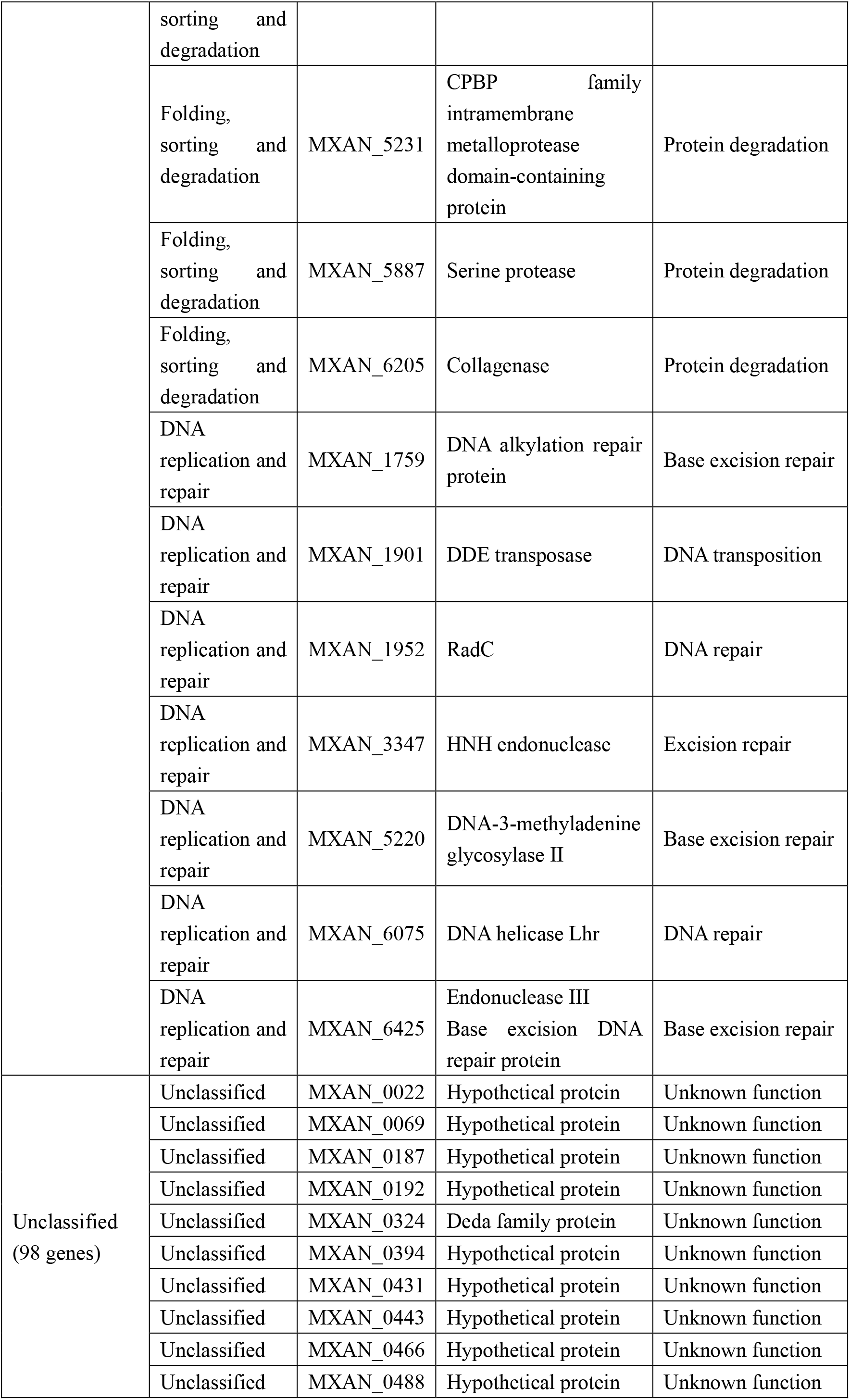
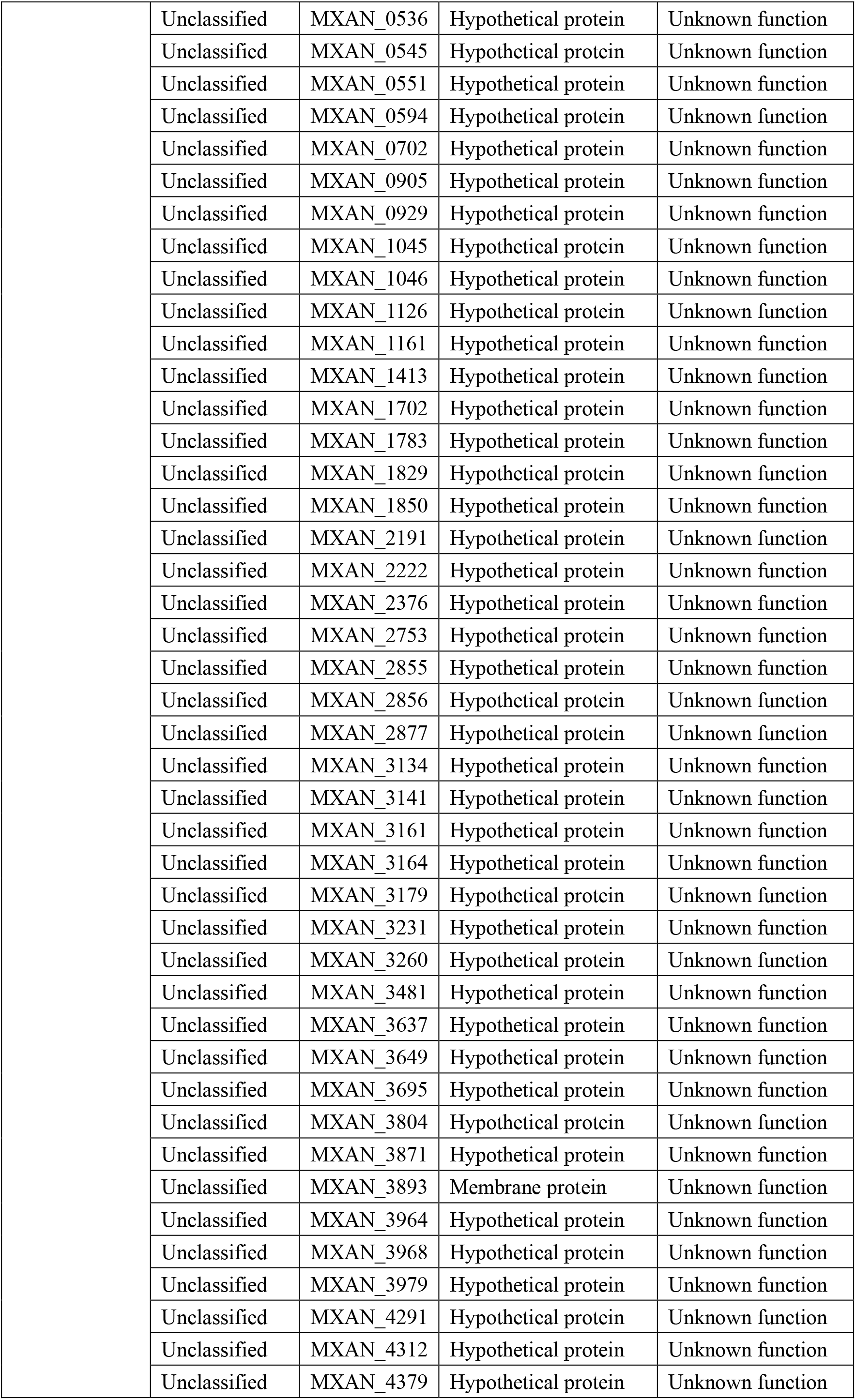
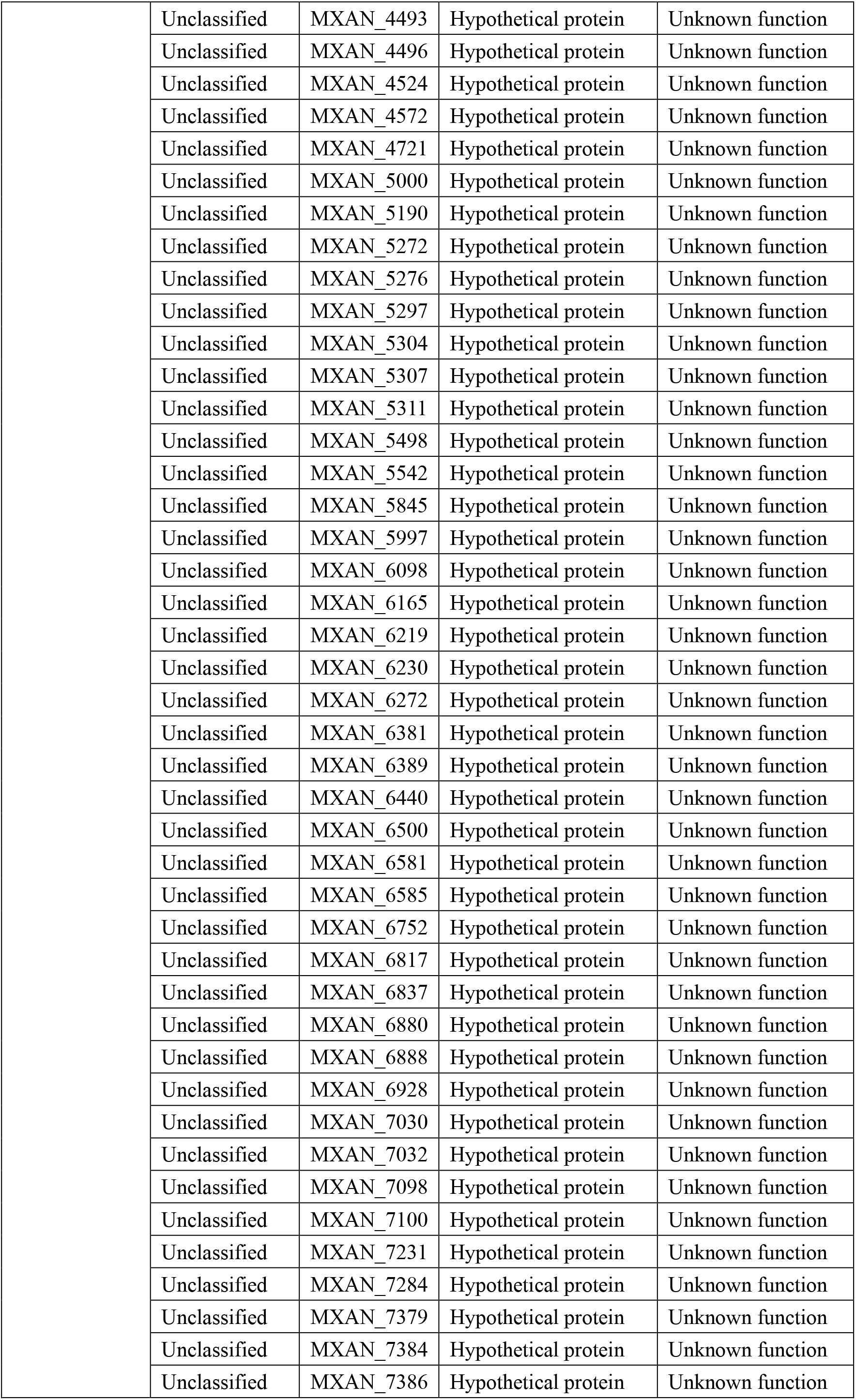
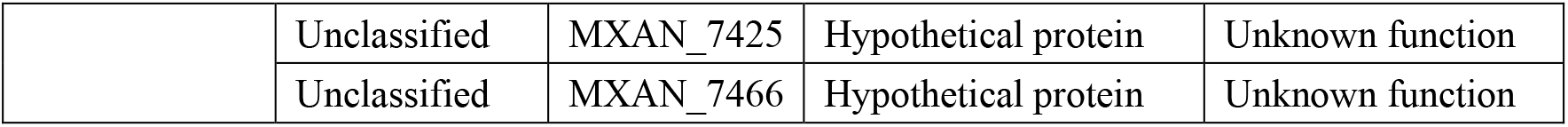
179 genes promoted by ImuA

**Table S3.**
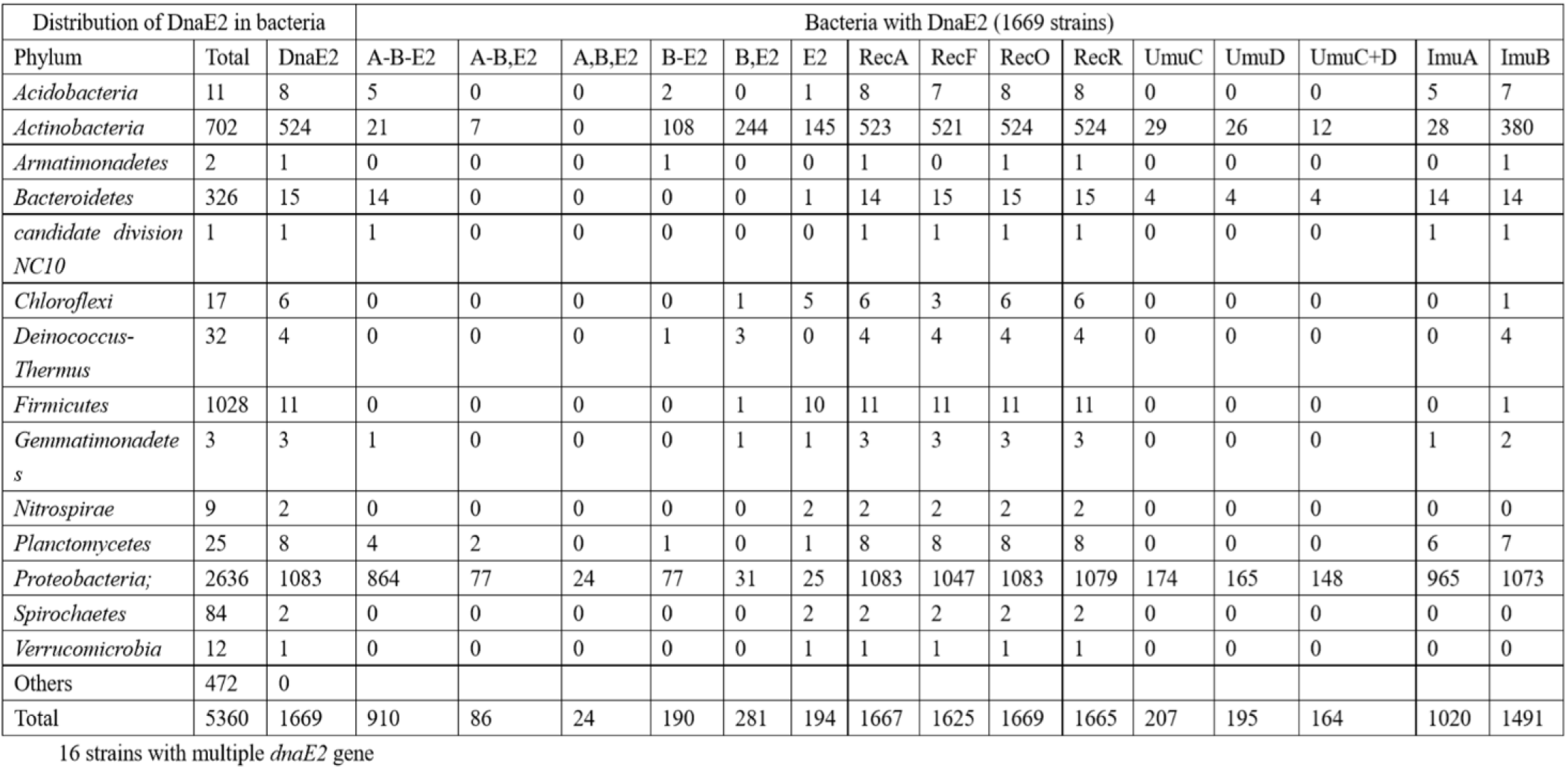
Appearance of genes related to DNA replication in sequenced bacterial genomes

**Table S4.** Strains and plasmids used in this study.

| Strains or plasmids | Genotype or description | Source or references |
| --- | --- | --- |
| **Strains** |  |  |
| *M. xanthus* |  |  |
| DK1622 | Wild-type strains | D. Kaiser, University of Stanford |
| IA | DK1622 ($\Delta imuA$) | This study |
| iA<sup>+</sup> | DK1622::pSW3991 (*imuA* with its own promoter integrated at *attB* site) | This study |
| IB | DK1622 ($\Delta imuB$) | This study |
| YL1601 | DK1622 ($\Delta dnaE2$) | Peng, 2017 |
| *E. coli* |  |  |
| DH5α(λpir) | F<sup>-</sup>80dlacZΔM15 $\Delta(lacZYA-argF)U169$ *deoR recA1 endA1 hsdR17* (*r<sub>k</sub><sup>-</sup>m<sub>k</sub><sup>+</sup>*) *phoA supE44 λ<sup>-</sup>thi-1 gyrA96 relA1* | Takara |
| BL21(DE3) | Expression strain, $\lambda$(DE3 [*lacI lacUV5-T7 gene 1 ind1 sam7 nin5*]) [*malB+*]K-12(λS) | This study |
| *S. cerevisiae* |  |  |
| AH109 | MATa, *trp1-901*, *leu2-3, 112*, *ura3-52*, *his3-200*, *gal4Δ*, *gal80Δ*, *LYS2::GAL1<sub>UAS</sub>-GAL1<sub>TATA</sub>-HIS3*, *MEL1 GAL2<sub>UAS</sub>-GAL2<sub>TATA</sub>-ADE2*, *URA3::MEL1<sub>UAS</sub>-MEL1<sub>TATA</sub>-lacZ* | Clontech |
| **Plasmids** |  |  |
| pBJ113 | Gene replacement vector with KG cassette, *Kam<sup>r</sup>* | Z.M. Yang, Virginia Tech |
| pBJ3991 | Upstream and downstream homologous arms of an *imuA* inserted into *EcoRI/HindIII* site of pBJ113, *Kam<sup>r</sup>* | This study |
| pGBK-T7 | Yeast two-hybrid vector, *Kam<sup>r</sup>* | Clontech |
| pGAD-T7 | Yeast two-hybrid vector, *Amp<sup>r</sup>* | Clontech |
| pSWU19 | Site-specific integration vector with Mx8 *attP* integration site, *Kam<sup>r</sup>* | (Wu & Kaiser, 1995) |
| pSW3991 | *ImuA* with its upstream 247bp promoter sequence, inserted into *XbaI/EcoRI* sites of pSWU19, *Kam<sup>r</sup>* | This study |
| pET3a | *E. coli* xpression vector, *Amp<sup>r</sup>* | Novagen |
| pET15b | *E. coli* xpression vector, *Amp<sup>r</sup>* | Novagen |

**Table S5.**
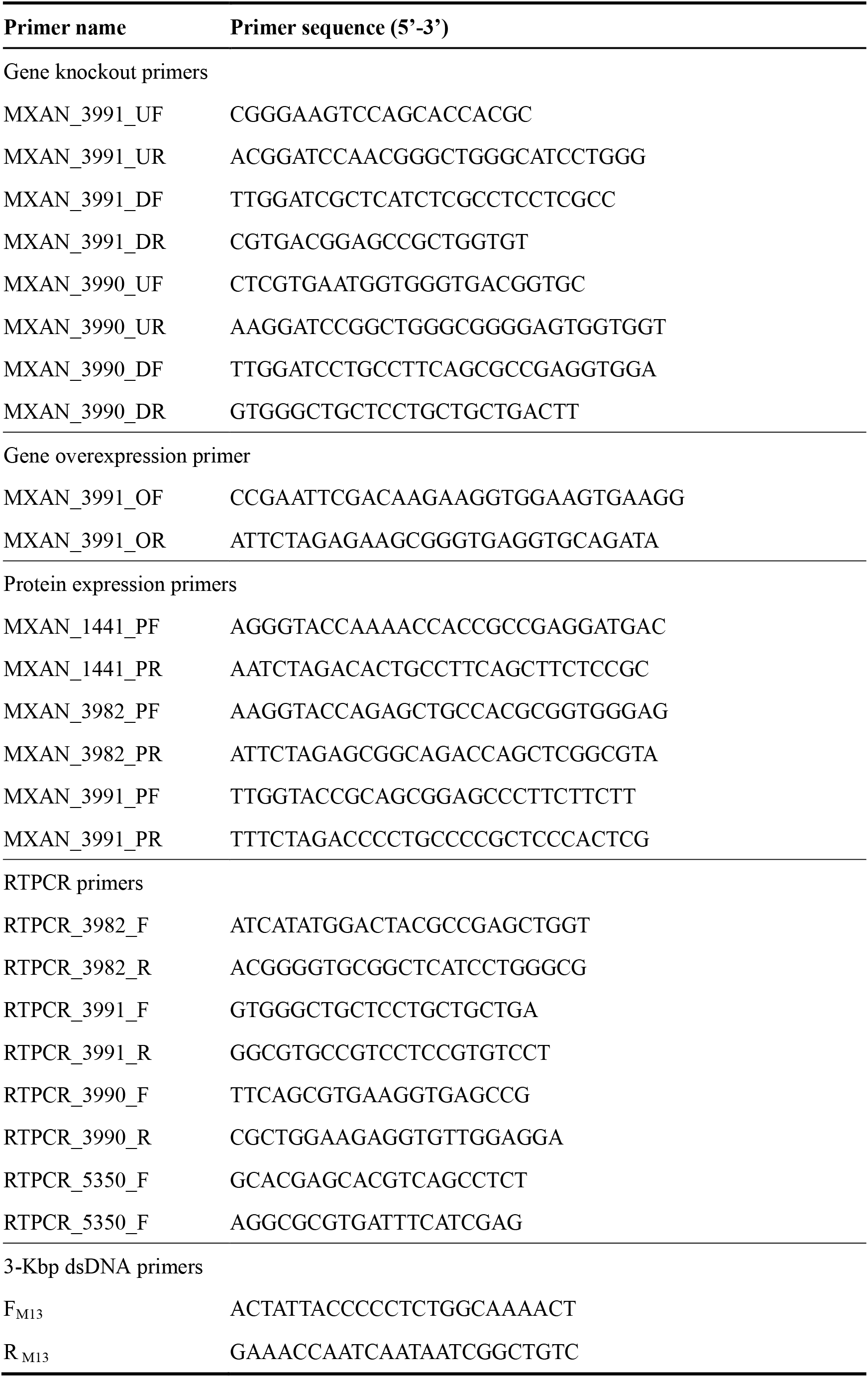
Primers used in this study.

